# Translating human drug use patterns into rat models: exploring spontaneous interindividual differences via refined drug self-administration procedures

**DOI:** 10.1101/2024.07.08.602268

**Authors:** Ginevra D’Ottavio, Sara Pezza, Jacopo Modoni, Ingrid Reverte, Claudia Marchetti, Soami F. Zenoni, Andrea Termine, Carlo Fabrizio, Daniela Maftei, Roberta Lattanzi, Giuseppe Esposito, Davide Ragozzino, Emiliano Merlo, Michele S. Milella, Roberto Ciccocioppo, Fabio Fumagalli, Marco Venniro, Aldo Badiani, Fernando Boix, Daniele Caprioli

**Affiliations:** Laboratory affiliated to Institute Pasteur Italia – Fondazione Cenci Bolognetti – Department of Physiology and Pharmacology, Sapienza University of Rome, Rome, Italy; Santa Lucia Foundation (IRCCS Fondazione Santa Lucia), Rome, Italy; School of Psychology, University of Sussex, Falmer, UK; Toxicology Unit, Policlinico Umberto I University Hospital, Rome, Italy; School of Pharmacy, Center for Neuroscience, Pharmacology Unit, University of Camerino, Camerino, Italy; Department of Pharmacological and Biomolecular Sciences, ‘Rodolfo Paoletti’, University of Milan, Milan, Italy; University of Maryland School of Medicine, Department of Anatomy and Neurobiology, Baltimore, USA; Section for Drug Abuse Research, Department of Forensic Sciences, Oslo University Hospital, Oslo, Norway

**Author notes:** Corresponding Author: Ginevra D’Ottavio Daniele Caprioli.

**Keywords:** heroin, pharmacokinetics, voluntary abstinence, incubation of craving, sex differences

## Abstract

Heroin and cocaine users tailor their dosage and frequency of use, as well as their method of administration, to maximize the drugs’ pleasurable effects and prevent withdrawal symptoms. On the other hand, many preclinical self-administration and choice experiments employ fixed unit doses and mandatory timeouts after doses (known as discrete dimension procedures). These restrictions fail to consider the distinct pharmacokinetic properties of heroin and cocaine, leading to uniform and comparable behaviors (including drug-taking patterns). This uniformity contrasts sharply with the significantly different ways humans use heroin and cocaine, which are characterized by highly individualized drug use behaviors. Here, we introduce a no-timeout procedure that overcomes this limitation (continuous dimension procedure).

We analyzed the heroin and cocaine taking- and seeking-patterns and estimated drug-brain levels in the presence or absence of timeout between drug injections. We further assessed how absence of timeout and the availability of drug or social peer (access time to the two rewards) affect drug preference. Removing the timeout had a profound effect on pattern of heroin taking and seeking, promoting the emergence of burst-like drug intake and social withdrawal as revealed by a discrete choice procedure. On the other hand, timeout removal had a lesser impact on cocaine taking and seeking and did not impact social preference. By removing timeout during self-administration and increasing the access time during choice resulted in a self-administration procedure that more closely mimic human heroin intake, offering a platform to identify novel medications.

## Introduction

Cocaine and heroin addictions persist as global health concerns, prompting ongoing research for novel treatments [1,2]. Preclinical research mainly contributes investigating the neurobiological effects of drug exposure by employing a plethora of animal models, with self-administration, along with its variants, as the ‘gold standard’ [3–6]. However, the extensive preclinical data amassed thus far has not resulted in ‘forward translation’ during the last two decades [3,6,7]. Determining the appropriate level of precision and the specific circumstances that warrant the refinement of such models is therefore central in the pursuit of this goal [6]. The present study was designed to help address this issue by using a behavior-centric approach. This approach suggest that “over-constrained” behavioral procedures may produce artificial behaviors and consequently potentially inaccurate neurobiological findings, with reduced translational power [8,9].

In the quest to establish a robust animal model, preclinical addiction neuroscientists leaned, and continue to lean, on an ‘across-drugs-convergent’ modeling strategy, based on the idea that drugs are positive reinforcers, similar to natural rewards [10–13]. This perspective was reinforced by the observation that most addictive drugs increase dopamine release [14,15]. In this framework, experimenters designed drug self-administration procedures with the goal to achieve regular drug-taking patterns comparable to natural rewards [10,16]. Notably, this approach also facilitates quick testing of hypotheses [17–19]. Regular patterns can be maintained by using “over-constrained” self-administration procedures characterized by implementation of discrete (as opposed to continuous) dimension strategies [18,20,21], involving experimenter-imposed unit-doses of drug interspersed by timeouts. Nevertheless, these strategies create a ‘unique case’ of drug-taking behavior that: 1) disregards drug metabolism, which often leads to the formation of active drug metabolites, contributing to behavioral effects [22–24]; 2) masks differences in patterns of drug taking between different addictive drugs; 3) prevents the experimental animal from self-selecting the appropriate dose-time relationship [18,21]; 4) masks interindividual differences in the pattern of self-administration [18] and in drug-versus-nondrug reward choice procedures [25], except when using specific experimentally-selected parameters [26–29]. Ideally, a self-administration procedure should encompass the behavioral clinical definition of the disorder itself and incorporate key dimensions of drug use (e.g., drug-taking patterns) [3,5,6,30]. The latter should emerge in the most spontaneous way as possible (i.e., least experimentally constrained) based on the unique pharmacological properties of the drug under investigation.

In real-world scenarios, individuals display the ability to instrumentalize drugs to achieve desired effects, responding to internal and external factors (e.g., drug availability, tolerance or sensitization to drug effects or side effects of excessive drug consumption) [23,31–37]. Heroin and cocaine users display markedly distinct drug instrumentalization. For example, they adopt distinct frequencies, dosages, and routes of administration [36–44]. These differences typically reflect pharmacokinetic and pharmacodynamic properties of the two drugs [23]. Notably, a relative common feature of cocaine and heroin users is transitioning from slower to faster routes of administration, with the aim of intensifying drugs’ effects [37,45–48]. This is particularly evident in a subset of individuals—often considered vulnerable—featuring excessive drug instrumentalization [referred to as over-instrumentalization [33]], leading to the development of severe and harmful patterns of use, culminating in substance use disorders [36,44,49–51].

Considering these aspects, here we advocate that the over-constraints typically implemented in drug self-administration and choice procedures might result in unnatural drug taking patterns and consequently influence other drug-related behaviors. To address this issue, we conducted a series of experiments, analyzing, and contrasting heroin and cocaine taking (fixed ratio and progressive ratio responding) and seeking (incubation of craving [52] patterns across various drug self-administration procedures, their estimated pharmacokinetics correlates [23,53,54], and their consequences on social peer-vs-drugs choice. We compared three self-administration procedure: 6-h continuous access [55] with or without 20-s timeout after injections, and 6-h intermittent access [56,57] without timeout. In the choice study, we also manipulated the duration of social access and drug exposure after each choice trial.

Overall, our study represents a step forward in refining animal models of addiction to be achieved by relaxing the constraints in self-administration experiments in order to reveal bona fide interindividual differences in animal models of addiction.

## Materials and methods

For a complete description of Materials and Methods and Statistical analysis see Supplementary Online Materials.

### Drug self-administration procedure

We carried out two experiments to investigate the impact of experimenter-imposed timeout between drug unit-doses on heroin and cocaine self-administration, using both within-subject and between-subject designs. To this aim we used an FR1 schedule of reinforcement with short injection times (3-s) and no timeouts between injections [strongly comparable to the hold-down drug self-administration procedure introduced by Morgan et al. [21]]. In experiment 1, we used a within-subject design to probe whether the timeout affects heroin and cocaine self-administration. We trained the same rats to self-administer drugs for 18 days with or without timeout using a two-lever drug self-administration procedure and we compared (1) patterns of drug taking; (2) drug-seeking under extinction conditions, (3) preference (in a discrete-choice procedure), and (4) motivation (using a recently developed progressive ratio procedure [18]) on the two access conditions.

In experiment 2, we explored the same research question using a between-subject design. We trained three groups of rats to self-administer heroin or cocaine for 15 days under three different self-administration training schedules: (1) continuous-access timeout (drugs continuously available 6-h/d, FR1 20-s timeout), (2) intermittent-access no-timeout (drugs intermittently available 6-h/d divided in 5-min access every 30-min, FR1 no-timeout), or (3) continuous-access no-timeout (drugs continuously available 6-h/d, FR1 no-timeout) schedule. Then, we compared rats‘ performance on: (1) overt behavior during self-administration [58,59], (2) patterns of drug taking, and resultant estimated drug brain levels during the last self-administration session, and (3) drug seeking under extinction conditions using the incubation of drug craving procedure [52].

### Drug-vs-social discrete choice procedure

In experiment 3, we tested whether the possibility to self-select the preferred dose in the preferred timing in the drug-vs-social choice would affect the preference between drug and social interaction. We trained rats for social self-administration and then for heroin or cocaine (drugs continuously available 6-h/d, FR1 no-timeout) and tested them for drug seeking. Then we tested rats’ preference for drug or social interaction in two different choice conditions: (1) in a choice procedure where rats are allowed to choose between 1 unit-dose of social interaction (1-min access) or drug (heroin dose 0.075 mg/kg/inf or cocaine 0.5 mg/kg/inf; (2) in a choice procedure where rats are allowed to choose between 5-min access to the social partner or to the drug (a time sufficient to self-administer the preferred dose in the preferred timing).

## Results

### Experiment 1. The role of timeout in drug self-administration: a within-subject study

The rats increased their heroin and cocaine intake over time in both access conditions (timeout and no-timeout) (**Figure 1D left, 1F left**). Heroin intake was notably higher without timeouts, while lever presses for heroin increased with timeouts. Notably, this effect was not observed with cocaine (**Figure 1D right, 1F right**). The rats exhibited stronger seeking for heroin and cocaine no-timeout (**Figure 2D, 2G**), strong preference for the no-timeout condition for both drugs during the choice tests (**Figure 2E, 2H**) and higher breaking points and cumulative number of lever presses for the no-timeout, relative to the timeout condition (**Figure 2F, 2I**).

**Fig. 1.**
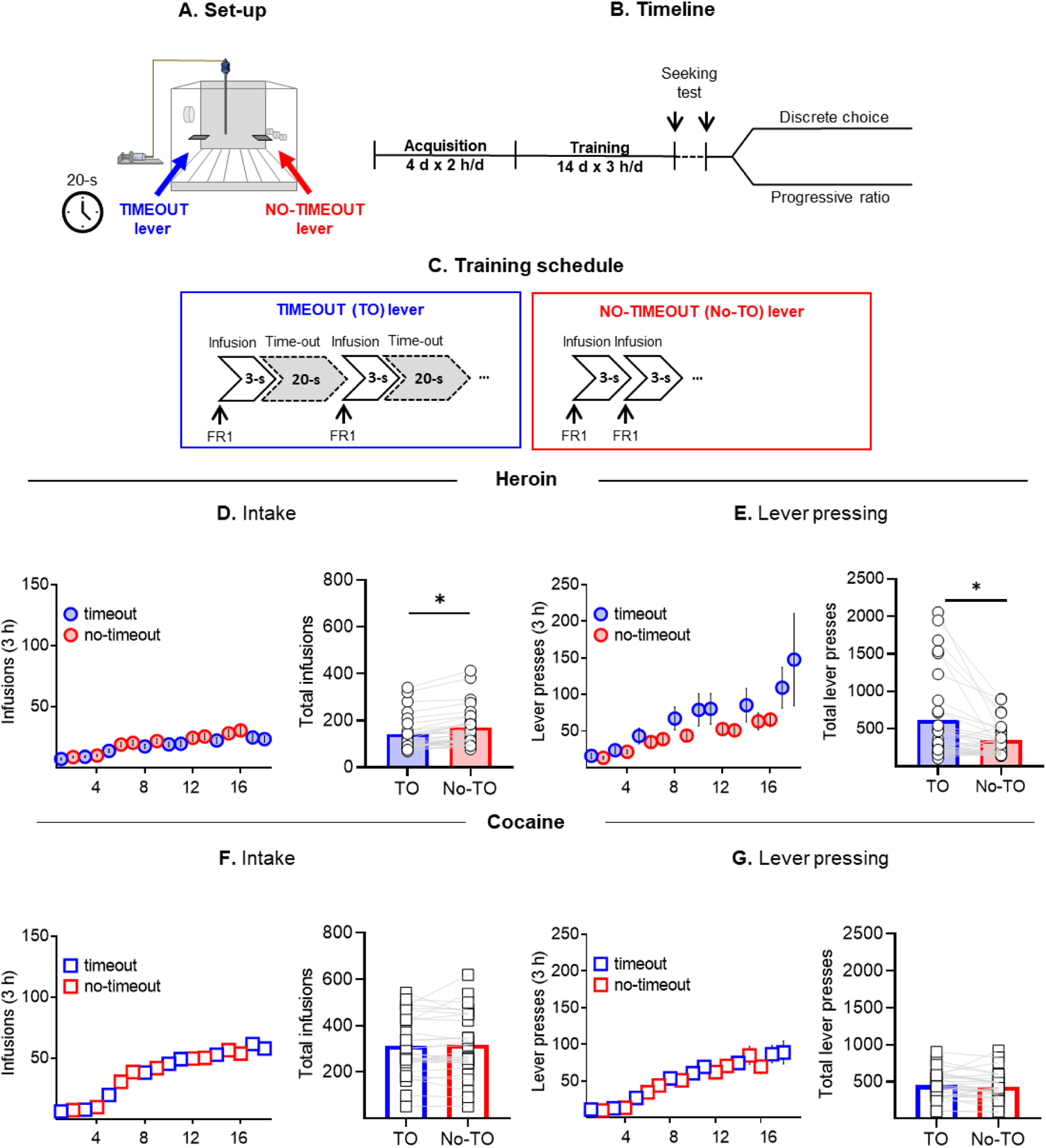
The impact of timeout on heroin and cocaine self-administration: comparison between timeout and no-timeout conditions: a within-subject design. (**A**) Experimental set-up. (**B**) Experimental timeline. (**C**) Training schedule. (**D and F**) Drug intake. Mean ± SEM number of drug infusions per session (left) and individual data of number of total drug infusions (right). (**E and G**) Lever pressing. Mean ± SEM number of lever presses per session (left) and individual data of number of total lever presses (right). *Different from timeout condition, p < 0.05 (heroin n = 29; cocaine n = 28).

**Fig. 2.**
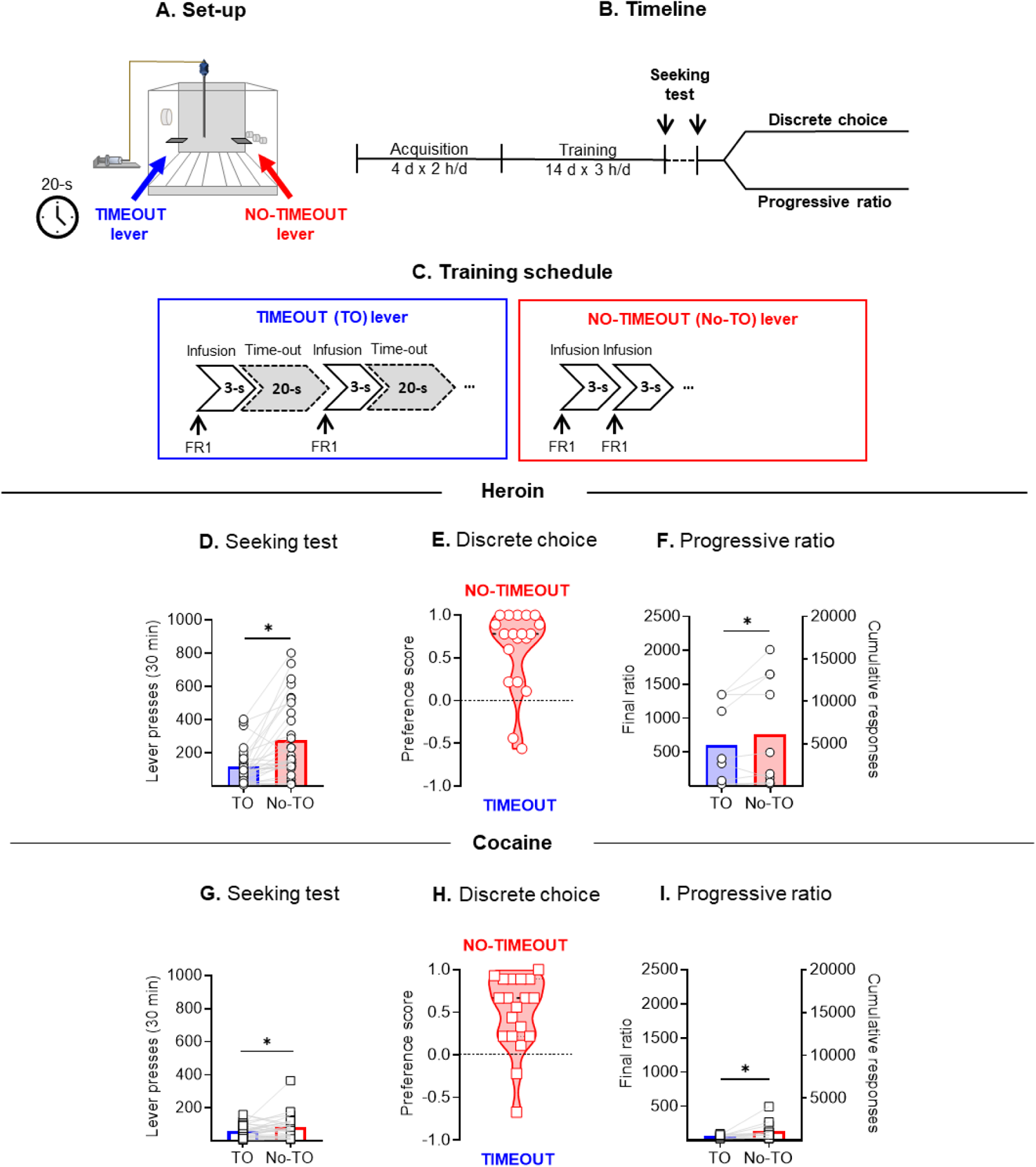
The impact of timeout on motivation to seek and take heroin and cocaine: comparison between timeout and no-timeout conditions in a within-subject design. (**A**) Experimental set-up. (**B**) Experimental timeline. (**C**) Training schedule. (**D and G**) Seeking test. Individual data of number of lever presses on the active lever during the 30-min extinction tests. (**E and H**) Discrete choice. Individual data of average preference score during the discrete choice sessions. (**F and I**) Progressive ratio. Individual data of final ratio completed and cumulative number of lever presses during the multi-day progressive ratio test. *Different from timeout condition, p < 0.05 (heroin n = 29; cocaine n = 28).

### Experiment 2. The role of timeout in heroin and cocaine self-administration: comparison between continuous-access timeout, intermittent-access, or continuous-access no-timeout

#### Drug intake

Across all access conditions (intermittent, continuous timeout, continuous no-timeout) drug intake and lever pressing increased over time (**Figure 3D, 3G**). However, we observed significant differences between the access conditions. Heroin intake was markedly higher in both intermittent and continuous-access no-timeout, relative to the timeout condition (**Figure 3E**). Interestingly, these differences in intake did not affect hyperalgesia development, though rats in the timeout condition experienced less heroin-related weight loss than those in other conditions (**Figure S1E, S1F**). Conversely, cocaine use was lower in intermittent access compared to continuous access (both timeout and no-timeout) conditions. However, total cocaine intake was higher in the continuous-access no-timeout relative to the continuous-access timeout condition (**Figure 3E, 3H**).

**Fig. 3.**
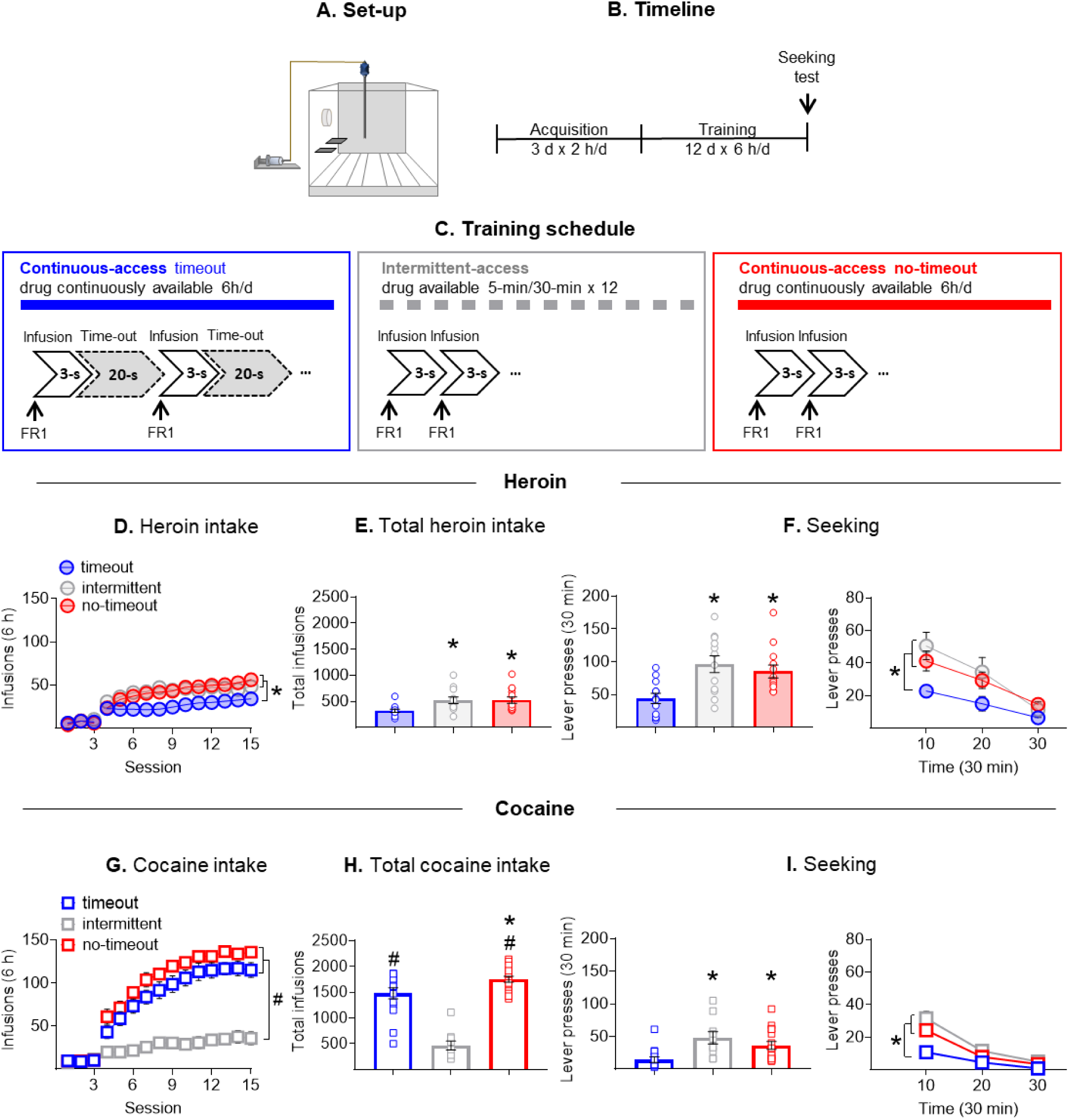
The impact of timeout in heroin and cocaine self-administration and seeking: a comparison between continuous-access timeout, intermittent-access, or continuous-access no-timeout to drug in a between-subject design. (**A**) Experimental set-up. (**B**) Experimental timeline. (**C**) Training schedule. (**D and G**) Drug intake. Mean ± SEM number of drug infusions per session. (E and H) Total heroin intake. Individual data of number of total drug infusions. (**F and I**) Seeking test on Abstinence day 1. Individual data of number of lever presses on the active lever during the 30-min extinction tests (left) and time course of lever presses extinction on the active lever during the 30-min extinction tests (right). Data are mean ± SEM of lever presses at each 10 min of the test session on day 1. * Different from continuous-access timeout, p < 0.05; # Different from intermittent-access, p < 0.05 (heroin n = 37; cocaine n = 41).

#### Drug-seeking

Overall, the rats displayed stronger seeking for heroin and cocaine self-administration without timeout (**Figure 3F, 3I**). We observed incubation of craving exclusively under timeout conditions, with increased lever pressing on day 21 compared to day 1 (**Figure S1D, S1G**).

#### Drug-taking patterns and estimated brain levels of drug

The different training conditions (**Figure 3D, 3G**) were associated with qualitative differences in the pattern of drug-taking (**Figure S2, S3**), with strong differences between heroin and cocaine. Regarding heroin, rats trained under continuous-access timeout conditions displayed a pattern of drug-taking characterized by few infusions (typically maximum 2 consecutive unit-doses) spaced by inter-infusion intervals of more than 10-min (**Figure S3D**). While without timeout (both intermittent and continuous access), rats consumed heroin in rapid, consecutive ‘bursts’ [60]: several infusions in a row, spaced by very short inter-infusions intervals (more than 2 consecutive injections in less than 5-min, **Figure S3D, S3E**). As in our previous study [51], ‘burst’ episodes were accompanied by fast-rising high brain peak concentrations of heroin and 6-MAM that were significantly higher in intermittent- and continuous no-timeout, relative to continuous timeout conditions (**Figure S4D, S4E**). While, morphine brain levels were significantly higher in continuous no-timeout, relative to continuous timeout and intermittent-access conditions (**Figure S4F**). Regarding cocaine, in both continuous timeout and no-timeout conditions, patterns of cocaine pattern followed a ‘loading’ then ‘maintenance’ pattern, leading to similar brain cocaine levels (**Figure 4G, 4H, 4I**), but ‘burst’ events were more frequent without timeouts (**Figure S3F**). In intermittent access, ‘burst’ episodes were separated by programmed time OFF (**Figure 4G**). Notably, this pattern was accompanied by brain cocaine peak concentrations and brain cocaine concentrations that were lower than in rats trained under continuous-access timeout and no-timeout conditions (**Figure 4H, 4I**).

**Fig. 4.**
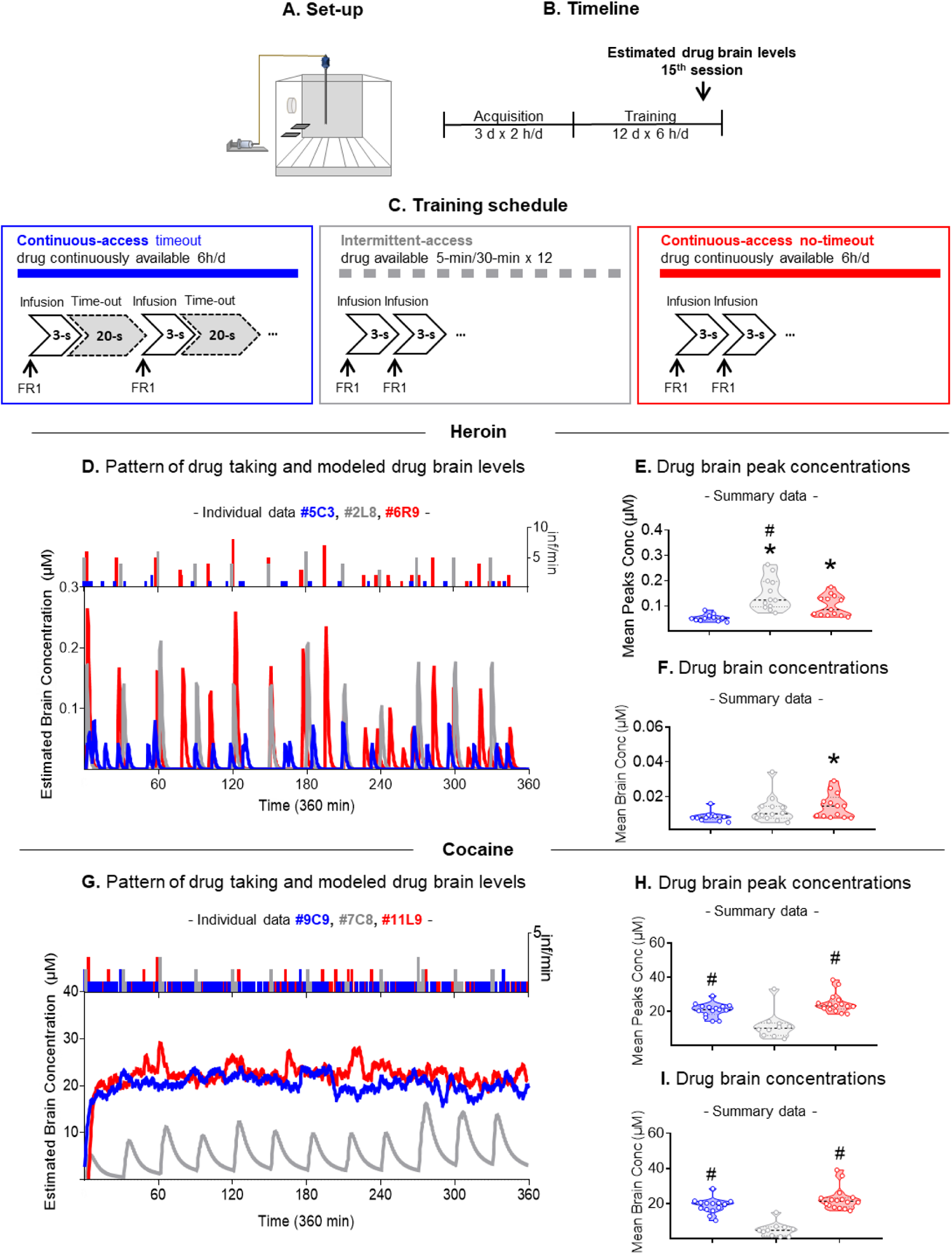
Patterns of drug intake and pharmacokinetic modelling of brain concentrations of heroin and cocaine in representative rats trained under continuous-access timeout, intermittent-access, or continuous-access no-timeout to drug in the last session of drug self-administration training. (**A**) Experimental set-up. (**B**) Experimental timeline. (**C**) Training schedule. (**D and G**) Pattern of drug-taking and estimated drug brain levels in representative rats (heroin infusions: continuous-access timeout = 30, intermittent-access = 55, continuous-access no-timeout = 86; cocaine infusions: continuous-access timeout = 137, intermittent-access = 44, continuous-access no-timeout = 146) (**E and H**) Estimated drug brain peak concentrations. Individual data of mean concentration of drug brain peaks. (**F and I**) Estimated drug brain concentrations Individual data of mean drug brain concentrations. Note the scales are adapted to the levels of each compound. * Different from continuous-access timeout, p < 0.05; # Different from intermittent-access, p < 0.05. (heroin: n = 37; cocaine: n = 41).

#### Behavioral repertoire

We observed that during the first hour of the last self-administration session heroin-trained rats showed similar behaviors across conditions. However, stupor was higher in intermittent and continuous no-timeout conditions, compared to continuous-access timeout (**Figure S6C**), but walking (locomotion) was higher in continuous-access timeout conditions (**Figure S6D**). In cocaine-trained rats walking was higher in intermittent conditions than in continuous-access timeout or no-timeout conditions (**Figure S6J**).

### Experiment 3. Discrete choice between drugs and social interaction

#### Discrete choice between drugs and social interaction in two different choice procedures

During the self-administration training, the rats increased their number of social and drug rewards earned over time (**Figure 5D, 5G**). After training, we divided rats into two groups, matched for drug intake (**Table 3, Supplementary Materials**): one group chose between 1 unit-dose of drug or 1-min (1 vs 1 unit-dose), while the other chose between 5-min of access to the drug or to the social partner (5 vs 5 min access). In the ‘1 vs 1’ choice, the heroin- and cocaine-trained rats strongly preferred social peer over drugs (**Figure 5F and 5I left**), as previously described [29,61]. In the ‘5 vs 5’ choice, cocaine-trained rats still preferred the social peer over higher drug doses (**Figure 5I**), and their preference did not change when the time of access to the drug was increased (**Figure S6F**). In contrast, we observed strong interindividual differences in preferences in heroin-trained rats (**Figure 5F right**): a subpopulation of heroin-trained rats preferred heroin over social interaction (**Figure 5F**) and their preference for heroin increased when the time of access to the social partner was increased (**Figure S7D**).

**Fig. 5.**
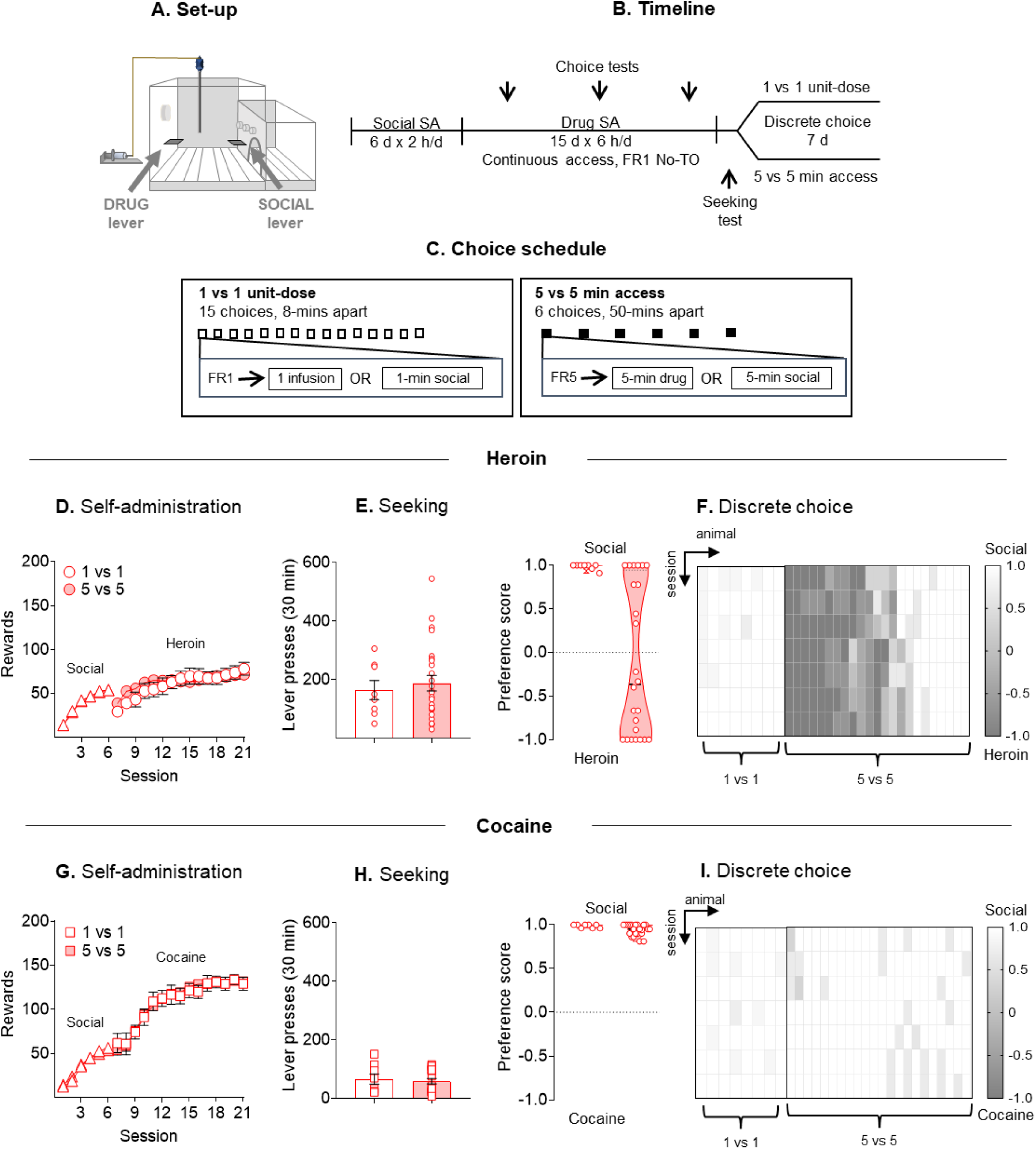
Discrete choice between drugs and social interaction in two different choice procedures. (**A**) Experimental set-up. (**B**) Experimental timeline. (**C**) Choice schedule. (**D and G**) Social and drug self-administration. Mean ± SEM number of social and drug rewards per session. (**E and H**) Seeking test. Individual data of number of lever presses on the active lever during the 30-min extinction tests. (**F and I**) Discrete choice. Individual data of mean preference score (left). Heatmap of individual social preference scores (right) across the 11 discrete choice sessions for rats tested in the two different choice procedures. White indicates a preference for social interaction (score = 1) and dark grey indicates a preference for drug (score = -1). (heroin: ‘1 vs 1 choice’ n = 8, ‘5 vs 5 choice’ n = 25; cocaine: ‘1 vs 1 choice’ n = 8, ‘5 vs 5 choice’ n = 24).

#### Interindividual differences in drug-vs-social choice: differences between heroin and cocaine

To understand the underlying factors influencing these preferences, we employed Principal Component Analysis (PCA) and Ascendant Hierarchical Clustering (AHC). PCA identified two main factors for each drug dataset (**Table S3**). For heroin, the two factors accounted for ∼71% of the total variance. The first factor (45.51%), included variables related to self-administration (Total Intake, Number of Bursts Events, and Peak Concentration) and Social Preference, while the second factor (26.07%), was solely associated with drug Seeking (**Table S3, Figure 6D**). Notably, despite a correlation between Seeking and Social Preference (**Figure 6C**), PCA indicated that variations in Social Preference were more significantly explained by drug-taking patterns. A negative correlation between Social Preference and the first factor suggested that more severe drug-taking patterns were linked to a lower preference for social interaction. Then we created a cumulative z-score (Severity z-Score) combining the three key self-administration variables (see above). z-scores negatively correlated with social preference (**Figure 7D**). Using an AHC analysis, we then categorized the rats into ‘vulnerable’ and ‘resilient’ clusters, starting from z-scores of the key self-administration variables (**Figure S9D**). Vulnerable rats had higher Severity z-scores (**Figure 7E**) and showed a stronger preference for heroin (**Figure 7F**), suggesting that the severity of drug self-administration could predict heroin-induced social withdrawal. In contrast, the cocaine dataset’s two factors explained ∼61% of the variance. The first factor included the self-administration variables and Seeking, while the second was solely related to Social Preference (**Table S3; Figure 7F**). This distinction implied that social preference variations were not directly linked to self-administration or seeking.

**Fig. 6.**
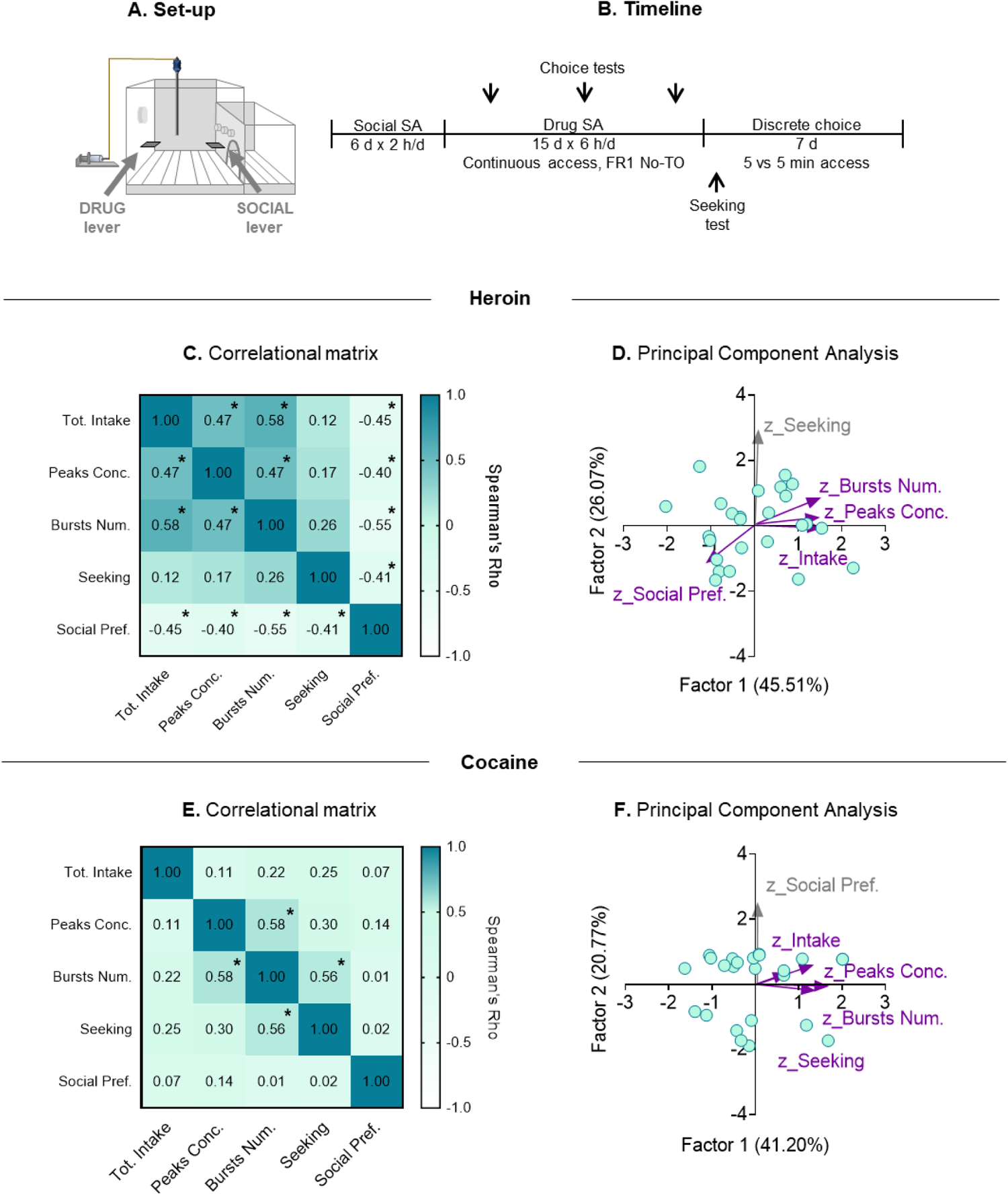
Interindividual variability in rats tested on the ‘5 vs 5 min’ choice procedure – Principal Component Analysis. (**A**) Experimental set-up. (**B**) Experimental timeline. (**C and E**) Correlational matrix. Correlational matrix of Spearman correlation between the main features of drug self-administrations (Total Intake, Number of Bursts Events, and Peaks Concentrations), Seeking and Social Preference (**D and F**) Principal Component Analysis. Biplot of the Principal Components Analysis (PCA) conducted on z-scores of the main features of drug self-administrations, drug seeking and social preference.

**Fig. 7.**
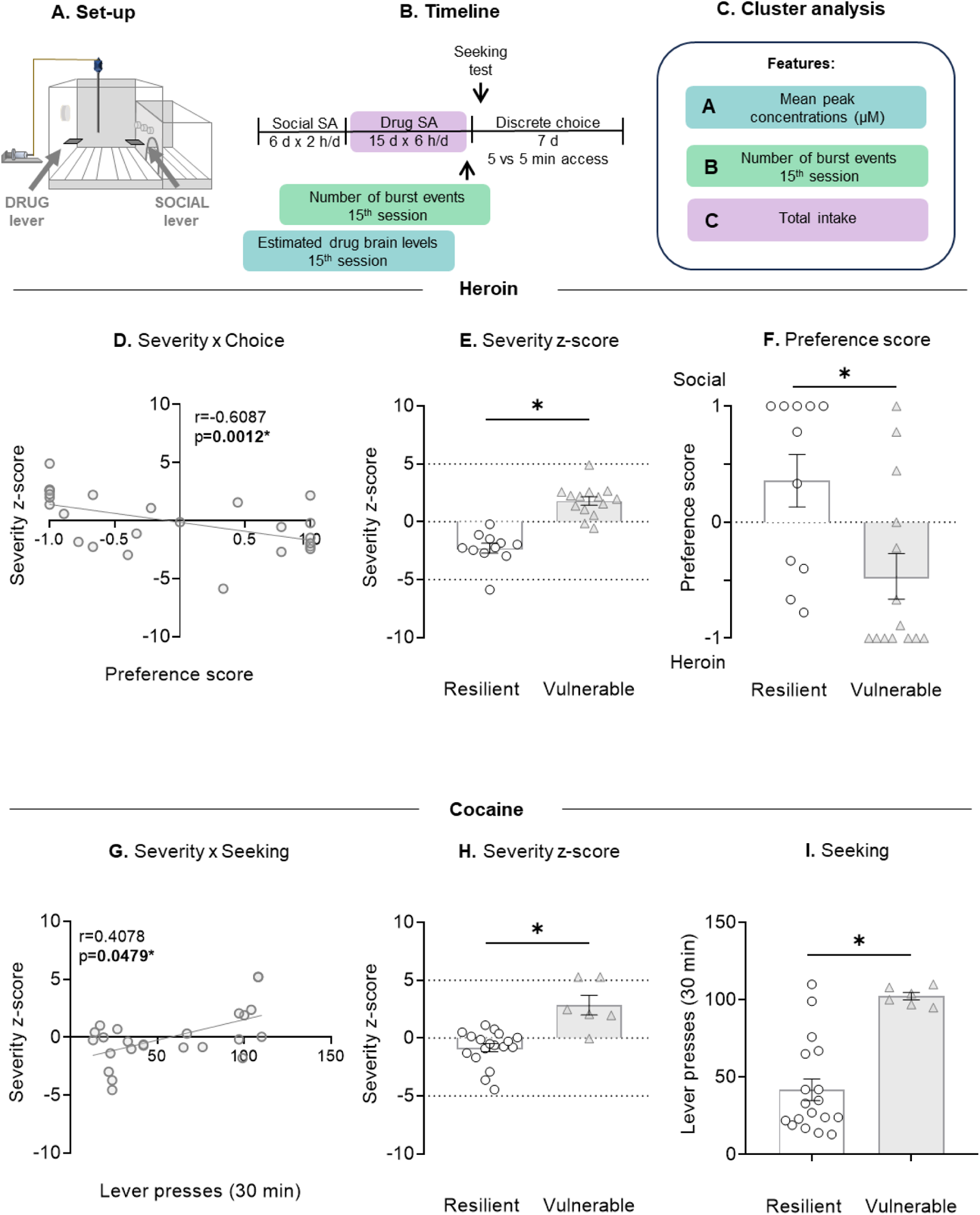
Addiction severity and interindividual variability in rats clusterized as vulnerable and resilient. (**A**) Experimental set-up. (**B**) Experimental timeline. (**C**) Features used for cluster analysis. (**D**) Correlation between Severity z-score and Choice. Correlation between Severity z score and Preference Score in heroin-trained rats. (**E and H**) Individual data of severity z-score in rats from resilient and vulnerable clusters. (**G**) Correlation between Severity z-score and Seeking. Correlation between Severity z score and Seeking in cocaine-trained rats. (**F**) Preference score. Individual data of mean social preference scores during the last 3 sessions of social choice in rats from resilient and vulnerable clusters. (**I**) Seeking. Individual data of drug seeking in rats from resilient and vulnerable clusters. * Different from resilient, p < 0.05. (heroin: n = 25; cocaine: n = 24)

However, a positive correlation between the Severity z-score and Seeking was observed (**Figure 7G**), indicating that more severe drug-taking patterns might explain increased seeking behavior (as previously observed [60]). AHC analysis for cocaine also identified ‘vulnerable’ and ‘resilient’ groups (**Figure S9E**), with vulnerable rats exhibiting higher Severity z-scores (**Figure 7H**) and more pronounced cocaine-seeking (**Figure 7I**).

## Discussion

We explored the effects of discrete versus continuous dimension strategies on drug self-administration and choice. Through a comparative analysis of cocaine and heroin, we found that implementation of timeouts and unit-doses: 1) decreases rats’ drive to seek and take drugs; 2) promotes standardized drug-taking patterns; 3) hinders the observation of spontaneous drug-taking patterns according to drugs’ pharmacokinetic profile, especially for drugs with short half-lives; 4) prevents the observation of relevant human drug-taking patterns; 5) prevents the emergence of interindividual differences in social vs. drug choice.

### The impact of timeout on drug self-administration

A key observation is that none of the rats trained without timeout experienced a fatal overdose. Initially, timeouts were used to create regular drug-taking patterns [10], but over time, the rationale for their implementation shifted to overdose prevention [62–65]. Here we showed that removing timeout, rather than leading to overdoses, results in non-standardized and spontaneous drug-taking patterns.

Under no-timeout conditions, rats were more likely to adopt bust-like drug-taking patterns [60]. These patterns were previously documented using intravenous cocaine and oxycodone self-administration with timeouts (from 10-s to 40-s) [60,66–68]. In our study, the occurrence of burst-like episodes of cocaine-taking was independent of timeout conditions; no-timeout only marginally increased the frequency of bursts events without affecting rats’ behavior. In contrast, burst-like patterns of heroin-taking under timeout conditions were not reported [54]. These patterns were observed exclusively using intermittent access to heroin, a procedure that do not incorporate timeouts [54,69]. However, here we observed that these patterns can appear under no-timeout conditions, regardless of access time to heroin (intermittent or continuous). In other words, the simple absence of timeout promotes burst-like heroin self-administration, typically resulting in ‘stupor’ (which is not observed in rats trained with timeout).

In our previous study, we speculated that timeout-related effects are linked to the drug’s metabolism: the faster the metabolism the stronger the impact of timeout on drug’s effects. Timeout between unit-doses prevents rats from self-administering drug at high frequency and, considering heroin’s brief half-life in the rat brain (∼0.9 min, [70]), this may hinder brain heroin accumulation, preventing rats from reaching high brain heroin concentrations and experiencing intoxication [22,54,70–76]. Here we further corroborated this hypothesis by using a simulation of brain concentrations of heroin, 6-MAM, and morphine in self-administration conditions with and without timeout. Simulated brain concentrations of heroin were halved in timeout, compared to the no-timeout conditions, while other metabolites were only limitedly affected.

In summary, while timeouts and specific experimental strategies influence heroin intake, they do not alter cocaine consumption [21] (**Figure 4**). This discrepancy is due to the unique pharmacokinetic and pharmacodynamic properties of each drug. Cocaine, has longer half-life in the rat brain—ranging from 8 to 11 minutes [77]—than heroin [70]. This suggests that timeout should have a minimal effect on brain cocaine accumulation. Accordingly, Minogianis et al. [78] demonstrated that varying the speed of cocaine delivery did not affect cocaine’s brain levels. Furthermore, studies on opioid drugs, like oxycodone – an opioid agonist like heroin but with a significantly longer half-life (60 minutes vs. heroin’s 0.9 minutes) [70,79,80] – reported burst-like pattern with timeout [67]. These findings indicate that timeout have a more pronounced impact on drugs with shorter half-lives.

Taken together, our study reinforces the notion that both pharmacokinetic and pharmacodynamic profiles of drugs are crucial in determining patterns of drug taking and seeking [23,81].

### Timeout strategies: bridging animal and human drug self-administration patterns

One of the aims of our study was to refine drug self-administration procedures in rats to mimic relevant drug-taking patterns observed in humans. In this regard, it is noteworthy that the *spontaneous* patterns of heroin-taking observed under no-timeout conditions closely resemble human taking patterns [82]. Indeed, individuals typically take large amounts of heroin, leading to intense euphoria, in a few daily episodes [31,35,41–43]. Based on previous studies, it has been argued that intermittent access procedures more closely mimic human behavior [53,57,83,84]. In contrast, our data suggest that continuous access to drugs, regardless of the presence of timeout, more closely resembles human drug-taking behaviors than intermittent access procedures. Indeed, individuals often engage in binges, consuming cocaine at very regular intervals [32,36–40,85]. From a refinement perspective, the significance our data is amplified in light of the FDA’s recent emphasis on drug use patterns as critical indicators of treatment success for SUDs [86–91], highlighting the necessity for animal models that accurately reflect human drug consumption to advance neurobiological research and treatment.

### The relationship between timeout, drug-taking patterns, and unit-doses

A main observation in our study is that rats did not limit themselves to the self-administration of single and spaced drug unit-doses. Despite considerable variability (**Figure S3**), when trained without timeout, rats displayed the tendency to self-administer multiple unit-doses in bursts. These findings challenge the idea that unit-doses *per se* correspond to the drug’s rewarding effects. Accordingly, Roberts and Zimmer [18], using a novel progressive ratio procedure, demonstrated that rats self-administering cocaine remain motivate to lever-press only if the dose of drug provided allow them to reach and maintain preferred drug brain levels. Notably, a single unit-dose of cocaine is not sufficient to reach these levels. We replicated these findings with cocaine [18] and generalized them to heroin. Overall, when rats were given access to their preferred drug dose, they self-administered multiple unit-doses of the drug and achieved high final ratios never documented in studies using progressive ratio schedules with single-unit doses [18,92–94].

Additionally, rats displayed higher motivation when their preferred dose was delivered faster, as observed by higher lever-pressing on the no-timeout lever. Accordingly, the speed of cocaine delivery strongly affects the motivation to take and seek the drug [95–100]. Additionally, while the specific influence of heroin delivery speed has not been fully explored, our data indicate that a faster heroin delivery intensifies heroin taking and seeking.

In this context it is important to note that, historically, the choice of doses and their timing in self-administration studies were often about convenience and standardization [101–103]. For example, studies justify dose reductions during self-administration training as a way to increase lever-pressing during self-administration and reinstatement tests [104–106]. However, our study challenges this viewpoint and suggests that the rewarding effects of drugs are closely linked to the administration of preferred doses at preferred time [18].

### The impact of timeout and unit-doses on drug-vs-social choice

Based on our self-administration results, we transitioned our investigation to drug-vs-social choice procedures. These procedures offer a way to mimic one of the hallmarks of drug addiction: greater behavioral allocation toward drugs over social-related rewards (family, employment) [30,107–119]. However, most preclinical studies on drug-vs-social choice procedures report strong preference for social interaction over intravenous drugs [29,61,120], with drug preference influenced by specific parameters (increased effort and delay of social reward) [26–29]. Nonetheless, in previous studies unit-dose strategies were used to evaluate choice behavior [120]. The use of unit-doses is typically justified based on the notion that they are on the descending limb of the drug dose-response curve [102,121,122]. Nevertheless, as previously described, unit-doses might not inherently determine the drugs’ reinforcing effects. Thus, we explore whether the provision of the preferred dose in the preferred timing in choice procedure affected drug preference.

Our data offered only partial validation to our hypothesis. We found that choice procedure variations predominantly influenced heroin, but not cocaine preference over social interaction. Overall, these data recapitulate previous studies demonstrating that social factors significantly influence drug self-administration [123]. Indeed, studies investigating effects of addictive drugs on direct social interaction indicate that opioids like morphine and heroin can lead to social deficits in rats and mice [124–133]. By contrast, research on cocaine’s effects on social behavior is limited and contradictory, with one study observing reduced social interaction [134] and another observing no significant change [130].

In the present study we found that heroin-induced social withdrawal (decreased preference for social interaction) can be observed exclusively in a subpopulation of rats and the social withdrawal can be predicted by the severity of drug-taking patterns. The more severe the drug-taking patterns, the greater the preference for heroin over social interaction. This observation is particularly relevant given the increasing focus on individual differences and factors influencing resilience and vulnerability to psychiatric conditions [49,135–138]. In this regard, our model could offer a potential framework for creating more effective, tailored interventions in human medicine.

Overall, our findings, together with previous research [130], suggest a dissociation on effects of opioids and psychostimulants on social behaviors in laboratory animals.

### Concluding remarks

Our study suggests a reevaluation of current methodologies used in preclinical drug self-administration models, such as the use of unit-doses and timeout, advocating for approaches that consider the unique pharmacological attributes of each addictive drug. Our results highlight the need to recognize the distinct characteristics of each drug, questioning the use of standardized self-administration protocols. By recognizing the complexity of drug effects and their interactions with behavioral patterns, we propose the use of models that address the multifaceted nature of addiction and its biological mechanisms, leading to more effective treatments.

## Acknowledgments

We thank Dr. Yavin Shaham for his comments on an earlier version of this manuscript and Miss Alana Sullivan for proofreading the manuscript. This work was supported by a grant from the “Enrico ed Enrica Sovena Foundation” in Rome, Italy, awarded for the fellowship of Mr. Soami Filippo Zenoni.

## Data availability statement

Materials, datasets, protocols and chamber details are available upon request to Ginevra D’Ottavio or Daniele Caprioli.

## Author contributions

Conceptualization: GD, DC, MSM, FB; Methodology: GD, DC, MSM, FB; Investigation: GD, SP, IR, CM, AT, CF, SFZ, IR; Visualization: GD, FB; Supervision: DC, MSM, FB; Writing—original draft: GD, DC; Writing—review & editing: AB, MV, MSM, FF, RC.

## Ethics approval statement

Procedures followed the guidelines of the national law (DL 26/2014) on the use of animals for research based on the European Communities Council Directive (2010/63/UE) and approved by the ethics committee of the Italian Ministry of Health and by local Ethical Committee of the Sapienza University of Rome.

