## Supplementary Online Materials for "Translating human drug use patterns into rat models: exploring spontaneous interindividual differences via refined drug self-administration procedures"

Ginevra D'Ottavio *et al.*

**This file includes:**

Supplementary Text

Figs. S1 to S10

Tables S1 to S9

References (24)

**Supplementary Text**

Subjects. A total of 282 male Sprague-Dawley rats (Charles River, Lecco) were used for the study. At the beginning of the experiments, the rats were 5-6 weeks old and weighed between 150-175 g. Before the surgery, the rats were pair-housed, and after the surgery, they were housed individually. For the social choice experiments, the rats were divided before the social self-administration training. Throughout the entire experiment, the rats were maintained on a reversed 12-hour light/dark cycle (lights off at 6 AM) with free access to standard laboratory chow and water. All the procedures followed the guidelines of national law (DL 26/2014) on the use of animals for research based on the European Communities Council Directive (2010/63/UE) and received approval from the ethics committee of the Italian Ministry of Health and the local Ethical Committee of the Santa Lucia Foundation. 19 rats were excluded due to catheter problems or sickness.

Drug. The drugs used in the study was donated from the Drug Supply Program of the National Institute on Drug Abuse (NIDA) and dissolved in sterile saline (0.9% NaCl). For drug self-administration training, unit-doses of 0.075 mg/kg/infusion for heroin and 0.5 mg/kg/infusion for cocaine were chosen based on previous studies [1,2].

Intravenous surgery. We anesthetized rats with isoflurane (5% induction, 2–3% maintenance) and injected them with Carprofen (2 mg/kg, subcutaneous injection; Zoetis Italia srl), immediately after the surgery and for the following 5 days to relieve pain and decrease inflammation. We inserted a silastic catheter into the jugular vein as previously described [3]. We placed the distal end of the catheter into the right jugular vein and attached the proximal end of a modified 22-gauge cannula placed on the back in the mid-scapular region. We flushed the catheters daily with a 0.2 ml sterile saline solution containing gentamicin (4.25 mg/ml; Fatro S.p.A.) to prevent occlusion during the recovery, training, and abstinence phases. We allowed rats to recover for a minimum of 5-7 days. We tested the catheter patency daily with the sterile saline and gentamicin solution and if during the training the catheter failed the test, we catheterized the left jugular vein, with the same procedure for the right, or we eliminated the rat from the study.

Self-administration apparatus. We trained rats in self-administration chambers placed inside sound-attenuating cubicles, which were equipped with an electric fan and controlled by a custom-made system.

The operant chambers had different components based on the specific experiment. The drug was delivered through a modified cannula (Plastics One; Roanoke, VA, USA) connected to a liquid swivel (Instech; Plymouth Meeting, PA, USA) via polyethylene-50 tubing that was protected by a metal spring.

Estimated brain levels of cocaine and heroin and its metabolites. To estimate the estimated brain levels of heroin and its active metabolites (6-MAM and morphine), and cocaine, we used the software program Kinetica v.5.1 (Thermo Fisher Scientific Inc., Waltham, MA, USA) with its FitMultiMicroExtravascular model for multiple administered doses. We based the estimations on the times of the single infusions during the last session of drug self-administration.

To estimate brain concentrations of heroin and its metabolites, we used the estimated pharmacokinetics parameters (absorption, elimination, intercompartmental distribution rate constants, time lag, volume of distribution) best fitting the mean brain extracellular fluid data for each opioid in the study by Gottås et al. [4] (see **Table 1**). To estimate brain concentrations of cocaine we used the pharmacokinetics parameters extracted from Pan et al. [5]. The brain concentrations were estimated from the absolute dose (µmol) received per infusion (adjusted to the weight of each animal) and were calculated in 0.5-min intervals for 8 hours, starting at the beginning of the session and up to 2 hours after its end. However, in the graphs, only the brain levels for the 6 hours of the session are presented.

*Experiment 1. The role of timeout in heroin and cocaine self-administration: a within-subject study*

Self-administration chambers. Chambers were equipped with a stainless-steel grid floor, and two operant panels were placed on the left and right walls. The left panel of the chamber was equipped with a house light and one active (retractable) lever. Responses on this lever activated the infusion pump and the discrete white-light cue located above the lever. The right panel was equipped with one active (retractable) lever. Responses on this lever activated the infusion pump and the three-light cue located above the lever. Left or right levers and relative cues were assigned randomly, to timeout or no-timeout conditions (see below).

Drug self-administration. Drug self-administration training was divided into two phases: acquisition and training. In the acquisition phase, we trained rats to self-administer drugs 2-h/d for 4d (acquisition phase; maximum infusions = 20/day). In the training phase, we trained rats 3-h/d for 14d (training phase; maximum infusions = 90/day). During both acquisition and training phase, we trained rats to self-administer drugs on two different levers: one lever associated with timeout and one lever associated with no-timeout. The timeout or no-timeout levers were randomly presented on alternate days. The sessions started with the insertion of one of the two levers (timeout OR no-timeout) and the illumination of the house light. Responses on the lever associated with the timeout condition (FR1) were reinforced by unit-doses of drug, paired with the cue light (3-s), followed by a 20-s timeout (cue light on), during which lever pressing was not reinforced and the cue light was on. Responses on the lever associated with the no-timeout condition (FR1) were reinforced by unit-doses of drug, paired with the cue light (3-s).

Drug self-administration. We tested rats for drug-seeking under extinction conditions on both levers (timeout and no-timeout) in the days following the last choice session. We tested rats on the two levers three days apart. The test days were balanced to avoid any confounding factors (abstinence days, lever-paired cues, etc.). The duration of the test sessions was 30-min to avoid a carryover effect on the second test. The sessions began with the illumination of the house-light, followed 10-s later by the insertion of the drug-paired lever; the house-light remained on for the duration of the session. Lever presses during the tests resulted in the contingent presentation of the light cue, previously paired with unit-doses of drug, but no drug infusion was delivered. After the last cue-induced seeking test, we divided rats (matched for their total drug intake during the self-administration training, Supplementary Table 3) into two groups. We tested one group of rats in a discrete choice procedure and the other group of rats was tested in a multi-day progressive ratio procedure.

Discrete-choice procedure. We tested a subgroup of rats for preference between timeout and no-timeout conditions in a discrete choice procedure where the rats were allowed to choose between the timeout-paired and no-timeout-paired levers. The discrete choice sessions lasted 6-h and were conducted using the same parameters (dose of drug and stimuli associated with the two levers) selected for the training phase. Each 6-h discrete choice session was divided into 6 trials separated by 55-min. The length of the inter-trial intervals was set based on a pilot study (data not shown) and corresponded to the time needed for morphine to dissipate [4]. Each trial began with the flashing of the house light for 30-s (a discriminative stimulus that signaled the choice session), followed by the fixed lighting of the house light and the insertion of both the timeout-paired and not-timeout-paired levers. Rats then had to select one of the two levers. The operant response requirement for the lever's selection was set to 5 consecutive responses (FR5) to avoid accidental choices. If the rats responded within 3-min, they had access to the selected lever for a total of 5-min. If the rats failed to respond on either active lever within 3-min, both levers were retracted, and the house light was turned off with no reward delivery. We tested rats starting from the day following the final drug self-administration session, continuing for a duration of 7 days.

Multi-day progressive ratio. We tested a subgroup of rats in a recently developed progressive ratio procedure [6], on both the timeout and the no-timeout levers, on alternate days in a random sequence. The progressive ratio procedure was identical to the training procedure except that access to the drugs was dependent upon an increasing number of lever presses (ratio value). The number of lever presses (ratio value) required was incremented within sessions and between days through the following progression: 1, 2, 4, 6, 9, 12, 15, 20, 25, 32, 40, 50, 62, 77, 95, 118, 145, 178, 219, 268, 328, 402, 492, 603, etc. [7,8]. Based on a recent study by Roberts and Zimmer [6], the completion of the ratio requirement provided access to 3-min to the drug under FR1 schedule. Sessions were 3h in length, after which time the levers were retracted. No restrictions were placed on the time required to complete the seeking ratio and no timeout periods were imposed after the 3-min access period. Each subsequent daily session started 1-step back to the ratio value reached on the day before. We excluded the rats if they did not exceed the final ratio value reached on the day before, corresponding to the breakpoint.

*Experiment 2. The role of timeout in drug self-administration: comparison between continuous-access timeout, intermittent-access, or continuous-access no-timeout to heroin and cocaine*

Self-administration chambers. Chambers were equipped with a stainless-steel grid floor, and one operant panel placed on the left wall. The panel of the chamber was equipped with a house light and the drug-paired active (retractable) lever. Responses on this lever activated the infusion pump and the discrete white-light cue located above the lever. In addition, the left panel was equipped with an inactive (stationary) lever that had no reinforced consequences.

Mechanical Nociceptive von Frey Test. We tested mechanical sensitivity with the von Frey Test only in rats that would then be trained for heroin, but not cocaine self-administration, because it was shown that mechanical sensitivity does not change after cocaine self-administration [9]. We tested rats before the drug self-administration training (7 days after the intravenous surgery) and after every three consecutive drug self-administration training sessions (in the morning, 12 hours into withdrawal) [9,10].

We put rats in boxes on an elevated metal mesh floor and allowed 10-min for habituation before examination. We stimulated the plantar surface of each hind paw with a series of von Frey hairs with logarithmically incrementing stiffness (0.04–2.0 g, 2Biological Instruments, Besozzo, Varese, Italy), presented perpendicular to the plantar surface (7–8 s for each hair). We determined The 50% paw withdrawal threshold (PWT) using Dixon's up-down method [11,12].

Drug self-administration. Drug self-administration training was divided into two phases: acquisition and training. In the acquisition phase, rats were trained to self-administer drug 2-h/d for 3d (acquisition phase; maximum infusions = 20/day). After the acquisition, we divided rats (matched for their total drug intake during acquisition, Statistical Table 2) into three groups that underwent three different drug self-administration training: continuous-access timeout, intermittent-access, and continuous-access no-timeout to drug. In the training phase, we trained rats 6-h/d for 12 days (training phase; maximum infusions: heroin = 90/d; cocaine = 150/d).

(1) Continuous-access timeout: drug continuously available 6-h/d; fixed ratio 1 (FR1) with 20-s timeout. Sessions started with the insertion of the two levers (active and inactive) and the illumination of the house light. Responses on the active lever (FR1) were reinforced by unit-doses of drug paired with the cue light (3-s), followed by a 20-s timeout during which lever pressing was not reinforced and the cue light was on.

(2) Intermittent-access: drug available in 12 epochs of 5-min (ON periods) every 25-min (OFF periods), FR1 no-timeout. Each 5-min ON period started with the insertion of the two levers and illumination of the house light and ended with the retraction of the levers and shutdown of the house light. Responses on the active lever were reinforced by unit-doses of drug, paired with a cue light (3-s), followed by no timeout.

(3) Continuous-access no-timeout: drug continuously available 6-h/d; FR1 no-timeout. Sessions started with the insertion of the two levers (active and inactive) and the illumination of the house light. Responses on the active lever (FR1) were reinforced by unit-doses of drug, paired with a cue light (3-s), followed by no timeout.

Behavioral repertoire assessment. We recorded rats' behavior during the first hour of the last drug self-administration training session. Sessions were recorded via a digital camera, videos were analyzed offline, and data were manually scored using Behavioral Observation Research Interactive Software [BORIS; [13]]. We scored the length of each bout of the following behaviors: grooming (corporal grooming and plucking/pulling at fur or digits), chewing, walking, stupor/immobility, hand/foot licking, and sniffing [14,15].

Seeking test. We tested rats for drug-seeking under extinction conditions on abstinence days 1 and 21. The duration of the test sessions was 30-min to minimize the carryover effect of extinction learning on day 1, which may decrease drug-seeking on day 21 [3]. The sessions began with the illumination of the house-light, followed 10-s later by the insertion of the drug-paired lever; the house-light remained on for the duration of the session. Lever presses during the tests resulted in the contingent presentation of the light cue, previously paired with unit-doses of drug, but no drug infusion was delivered. After the relapse test on day 1, we brought the rats to their home cages and handled them twice a week during abstinence.

*Experiment 3. Drug versus social interaction choice*

Self-administration chambers. Chambers were equipped with a stainless-steel grid floor, and two operant panels were placed on the left and right walls. The left panel was equipped with a house light and the drug-paired active (retractable) lever. Responses on this lever activated the infusion pump and the discrete white-light cue located above the lever. The right panel was equipped with the social partner-paired active (retractable) lever. Responses on this lever activated the three-light cue located above the lever and determined the opening of the guillotine-style sliding door.

Social self-administration. The training procedure was similar to the one described in previous studies [16]. We trained rats to press a lever to have access to a social partner 2-h/d for 6d. The resident rats were housed with their social partner (cage mate) until 7 days prior to social interaction self-administration, and each resident rat lever pressed for its previously paired partner. The training sessions started with the illumination of the house light and the insertion of the social partner-paired lever (that remained inserted throughout the session); responses on this lever resulted in access to the social partner (1-min), paired with the illumination of the three-light cue (1-min).

Drug self-administration. After the social self-administration, we trained rats to self-administer drugs 6-h/d for 15 days under continuous-access no-timeout conditions. Drug self-administration training sessions started with the insertion of the lever and the illumination of the house light. Responses on the active lever (FR1) were reinforced by unit-doses of drug (3-s), paired with the cue light (3-s). After every three consecutive drug self-administration sessions, we tested rats for drug versus social interaction preference in a discrete choice procedure (see below). We randomly assigned rats to the two choice conditions.

Seeking test. We tested rats for drug-seeking under extinction conditions on abstinence day 1. The duration of the test sessions was 30-min. The sessions began with the illumination of the house-light, followed 10-s later by the insertion of the drug-paired lever; the house-light remained on for the duration of the session. Lever presses during the tests resulted in the contingent presentation of the light cue, previously paired with unit-doses of drug, but no drug infusion was delivered. After the relapse test on day 1, we brought the rats to their home cages.

Discrete trial choice. We conducted the choice procedure using the same parameters (dose of drug and time of access to the social partner per reward and stimuli associated with the two active retractable levers) selected for the training phase. We allowed rats to choose between the drug-paired and the social partner-paired levers. We tested rats for a total of 7 days.

(1) Discrete choice '1 vs 1 unit-dose'. Choice sessions lasted for 120-min. Each 120-min choice session was divided into 15 discrete trials that were separated by 8-min. Each trial began with the presentation of the house light followed 10-s later by the insertion of both the social partner-paired and drug-paired levers. Rats then had to select one of two levers. The operant response requirement for the lever's selection was set to two consecutive responses (FR2) to avoid accidental choices. If the rats responded within 3-min, they received the reward corresponding with the selected lever. Reward delivery was signaled by the social partner-paired cue (1-min) or drug-paired cue (20-s), the retraction of both levers and the turning off of the house light. If the rat failed to respond on either active lever within 3-min, both levers retracted, and the house light was turned off with no reward delivery [16].

(2) Discrete choice '5 vs 5 min access'. Choice sessions lasted for 6-h. Each 6-h choice session was divided into 6 discrete trials that were separated by 55-min. Each trial began with the flashing of the house light for 30-s (a discriminative stimulus that signaled the choice session), followed by the illumination of the house light and the insertion of both the social partner-paired and drug-paired levers. Rats then had to select one of the two levers. The operant response requirement for the lever's selection

was set to 5 consecutive responses (FR5) to avoid accidental choices. If the rats responded within 3-min, they had access to the selected lever for a total of 5-min under FR1 schedule. The duration of access to the social partner was matched with the time allocated to drug access. Notably, Chow et al. [17] demonstrated that the time of access to the social partner does not impact progressive ratio responding. If the rats failed to respond on either active lever within 3-min, both levers were retracted, and the house light was turned off with no reward delivery.

<u>Discrete choice with incremental access time to drug or social partner.</u> We tested a subgroup of rats for 4 consecutive days in a choice procedure with incremental access time to drug (in the case of cocaine) or social partner (in the case of heroin). To allow rats to evaluate each option separately before expressing their choice, each choice session was preceded by 4 sampling trials spaced by 55-min. During the sampling, each trial began with the flashing of the house light for 30-s (discriminative stimulus that signaled the session), followed by the fixed illumination of the house light and the insertion of only one of the two levers (drug-paired, social-paired) in the following order: drug – social – drug – social. We counter-balanced the order of lever presentation (i.e., whether drug- or social-paired lever was presented first) across rats. During the sampling, rats were required to complete each trial to advance [18]. During the choice (4 trials), each trial began with the flashing of the house light for 30-s (discriminative stimulus that signaled the session), followed by the illumination of the house light and the insertion of both the social partner-paired and drug-paired levers. During sampling and choice, the response requirement was set to 5 (FR5) consecutive responses. We tested rats trained to self-administer heroin for preference between drug and social interaction in a choice procedure where the access time to the social partner was randomly increased across testing days from 1-min to 15-min and the access time to heroin was maintained constant to 1-min (see **Figure S7C**). On the contrary, we tested rats trained to self-administer cocaine for preference between drug and social interaction in a choice procedure where the access time to cocaine was randomly increased across testing days from 1-min to 15-min and the access time to the social partner was maintained constant to 1-min (see **Figure S7E**).

<u>Discrete-choice trial choice on late abstinence.</u> On day 60 from the beginning of the experiment, we tested rats for discrete choice after prolonged abstinence. This choice test was preceded by 4 sampling trials, as described above, to allow rats to evaluate each option separately before expressing their choice.

After the sampling, we allowed the rats to choose between the social partner-paired and drug-paired levers. During the choice (4 trials), each trial began with the flashing of the house light for 30-s (discriminative stimulus that signaled the session), followed by the illumination of the house light and the insertion of both the social partner-paired and drug-paired levers.

*Statistical analysis*

Experiment 1. For the training phase, we analyzed total drug intake and total lever presses for infusions/number of lever presses using the within-subjects factor of Session. We analyzed drug intake and lever presses during self-administration training using a GLMM as a function of Access (timeout, no-timeout) condition. In the GLMM, we used the Access condition as a fixed effect, while we used Sessions and Rat as random effects in a crossed design. For the relapse test, we analyzed active lever presses during the cue-induced seeking tests using the Wilcoxon matched-pairs signed-rank test with Access condition (timeout, no-timeout) as within-subjects factor; we included Cues (white light or three-light cue) and Abstinence Day (1,3) as covariates. Relative to the choice procedure, we calculated the preference (preference score) in the choice tests by normalizing the indifference level between timeout and no-timeout choices at 0 using the following formula: [1 - (% timeout lever choices/50%)] [19]. We analyzed the preference score across sessions using the within-subjects factor of Session. Finally, for the progressive ratio test, we analyzed the total number of lever presses using the Access condition (timeout, no-timeout) as within-subjects factor.

Experiment 2. For the training phase, we analyzed drug intake and lever pressing data for infusions/number of lever presses using the between-subject factor of Access condition continuous-access timeout, intermittent-access, and continuous-access no-timeout) and the within-subjects factor of Session. Regarding the pattern of drug self-administration and pharmacokinetics data, estimated based on behavioral data collected during the last session of drug self-administration, we analyzed total drug intake of the last session of self-administration, number of bursts-like events (defined as ≥3 infusions in less than 5-min, similarly to what previously described by Belin et al. [20]), number of infusions per peak, mean drug brain peak concentrations (calculated from valley to maxima), mean drug brain concentration and mean drug brain peak concentrations slope, using the between-subject factor of Access (continuous-access timeout, intermittent-access, and continuous-access no-timeout). Concerning the behavioral observations, collected during the last self-administration session, we analyzed each behavior (Stupor, Walking, Chewing, Grooming, Sniffing, Hand licking) using the between-subject factor of Access (continuous-access timeout, intermittent-access, and continuous-access no-timeout). Regarding the relapse test on abstinence day 1, we analyzed the number of active-lever presses using the between-subject factor of Access (continuous-access timeout, intermittent-access, and continuous-access no-timeout). We compared drug-seeking on Abstinence day 1 and 21 by analyzing the number of active lever presses using the between-subject factor of Access (continuous-access timeout, intermittent-access, and continuous-access no-timeout) and the within-subjects factor of Abstinence day (1, 21); the inactive lever was included as a covariate. Then, we analyzed withdrawal signs (hyperalgesia, body weight) using the between-subject factor of Access (continuous-access timeout, intermittent-access, and continuous-access no-timeout) and the within-subjects factor of Time (baseline, test days).

Experiment 3. For the training phase, we analyzed drug intake and lever presses for infusions/number of lever presses using the between-subject factor of Choice condition (1 vs 1, 5 vs 5) and the within-subjects factor of Session. Relative to the choice procedure, we calculated the preference (preference score) in the choice tests by normalizing the indifference level between social and drug choices at 0 using the following formula: [1 - (% social choices/50%)]. Then, we analyzed the preference score across sessions using the between-subject factor of Choice condition (1 vs 1, 5 vs 5) and the within-subjects factor of Session. Relative to the relapse test, we analyzed the number of active-lever presses using the between-subjects factor of Choice condition (1 vs 1, 5 vs 5). We then analyzed individual variability in the group of rats testes on the 5 vs 5 Choice procedure. To this aim we conducted Principal Component Analysis (PCA) and then Ascending Hierarchical Clustering (AHC). We normalized the data using z-scoring. To calculate z-scores we used the following formula: $z = (x-\mu)/\sigma$, where x is the individual value for the variable of interest, μ and σ the mean and standard deviation of the dataset for the variable of interest. We performed z-score normalization for five variables of interest: the main features of drug self-administration Total Intake, Number of Bursts Events, and Peaks Concentrations; Seeking and Preference Score. After normalization we used the Bartlett’s test of sphericity and the Kaiser-Meyer-Olkin (KMO) measure of sampling adequacy to determine whether our dataset was adequate for dimensional analysis. Subsequently, we conducted PCA (XLSTAT) to examine the underlying dimensionality of the normalized variables. We then investigated whether the severity of drug intake and patterns of drug-taking are connected to or could predict other addiction-like behaviors. To this aim, we aggregated z scores of main features of drug self-administration in a Severity z-score index and correlated it with Seeking and Social Preference. Finally, we performed a cluster analysis, as previously described [21-24]; to examine if variations in drug self-administration patterns and brain drug levels could delineate distinct subgroups, each displaying differences in preferences in drug Seeking or Social Preference. We used AHC (XLSTAT) (Ward's method) to find optimal clusters classification of mice into two clusters, based on z-scores. We used this methos since it does not use a target field. For clustering, we used main features of drug self-administration: (1) Total Intake, (2) Number of Bursts Events and (3) Peaks Concentrations. After clustering, we examined whether the clusters selected could predict the magnitude of Seeking or Social Preference. We analyzed the Seeking or Preference Scores (average over the last three choice sessions – a period where preference was stable) utilizing the between-subject factor of Cluster (vulnerable, resilient).

*Fig. S1. The impact of timeout in heroin and cocaine self-administration: craving and withdrawal signs after continuous-access timeout, intermittent-access, or continuous-access no-timeout to drug.*

(**A**) Experimental set-up. (**B**) Experimental timeline. (**C**) Training schedule. (**D** and **G**) Seeking test on Abstinence day 1 and 21. Individual data of number of lever presses on the active lever during the 30-min extinction tests. (**E** and **H**) Body weight. Mean ± SEM of body weight per session. (**F**) Von Frey test (hyperalgesia). Mean ± SEM of % change from baseline in paw withdrawal per session. *Different from day 1, $p < 0.05$ (heroin: n=37; cocaine: n=41).

**Figure S1.**

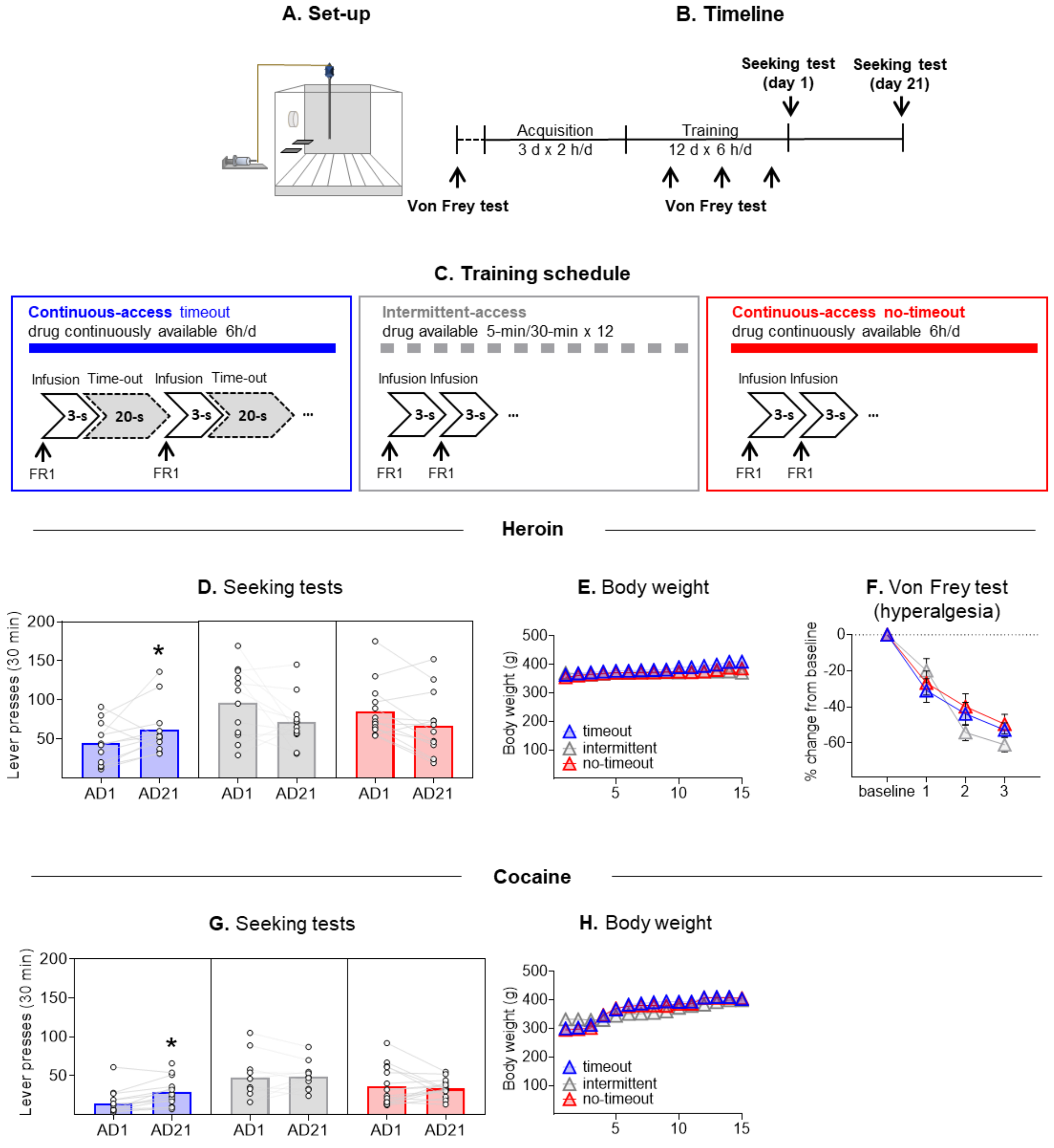

A. Set-up
B. Timeline
Seeking test (day 1)
Seeking test (day 21)
Acquisition
3 d x 2 h/d
Training
12 d x 6 h/d
Von Frey test
Von Frey test
C. Training schedule
Continuous-access timeout
drug continuously available 6h/d
Infusion
Time-out
Infusion
Time-out
3-s
20-s
3-s
20-s
FR1
FR1
Intermittent-access
drug available 5-min/30-min x 12
Infusion Infusion
3-s
3-s
FR1
FR1
Continuous-access no-timeout
drug continuously available 6h/d
Infusion Infusion
3-s
3-s
FR1
FR1
Heroin
D. Seeking tests
Lever presses (30 min)
200
150
100
50
*
AD1
AD21
AD1
AD21
AD1
AD21
E. Body weight
Body weight (g)
500
400
300
200
100
timeout
intermittent
no-timeout
5
10
15
F. Von Frey test (hyperalgesia)
% change from baseline
0
-20
-40
-60
baseline
1
2
3
Cocaine
G. Seeking tests
Lever presses (30 min)
200
150
100
50
*
AD1
AD21
AD1
AD21
AD1
AD21
H. Body weight
Body weight (g)
500
400
300
200
100
timeout
intermittent
no-timeout
5
10
15

*Fig. S2. The impact of timeout in heroin and cocaine self-administration: patterns of drug-taking on the last drug self-administration session.*

(**A**) Experimental set-up. (**B**) Experimental timeline. (**C**) Training schedule. Mean (**D** and **F**) and individual (**E** and **G**) cumulative infusions in the last self-administration session. (heroin: n = 37; cocaine: n = 41).

**Figure S2.**

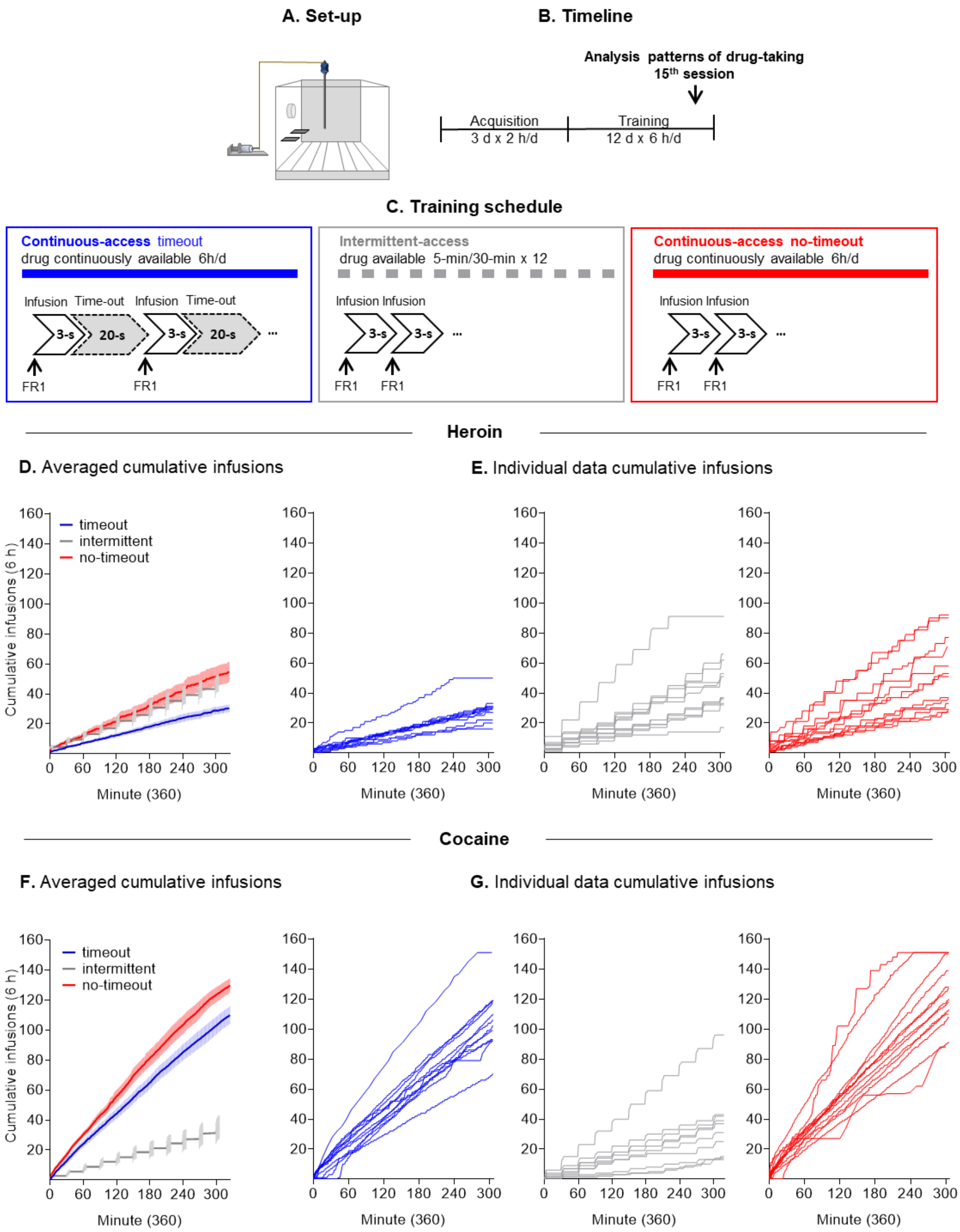
A. Set-up
B. Timeline
Analysis patterns of drug-taking
15th session
Acquisition
3 d x 2 h/d
Training
12 d x 6 h/d
C. Training schedule
Continuous-access timeout
drug continuously available 6h/d
Infusion Time-out Infusion Time-out
3-s 20-s 3-s 20-s ...
FR1 FR1
Intermittent-access
drug available 5-min/30-min x 12
Infusion Infusion
3-s 3-s ...
FR1 FR1
Continuous-access no-timeout
drug continuously available 6h/d
Infusion Infusion
3-s 3-s ...
FR1 FR1
Heroin
D. Averaged cumulative infusions
E. Individual data cumulative infusions
timeout
intermittent
no-timeout
Cumulative infusions (6 h)
Minute (360)
Cocaine
F. Averaged cumulative infusions
G. Individual data cumulative infusions
timeout
intermittent
no-timeout
Cumulative infusions (6 h)
Minute (360)

*Fig. S3. The impact of timeout in heroin and cocaine self-administration: frequencies of mean inter-infusion intervals and mean number of consecutive infusions per 2-min periods in continuous-access timeout, intermittent-access, or continuous-access no-timeout to drug.*

(**A**) Experimental set-up. (**B**) Experimental timeline. (**C**) Schematic definition of burst-like event. (**D**) Training schedule. (**E and G**) Frequency of inter-infusion intervals. (**F and H**) Number of burst-like events. Mean ± SEM of number of events. *Different from continuous-access timeout, $p < 0.05$; #Different from intermittent and continuous-access timeout (heroin: n=37; cocaine: n=41).

**Figure S3.**

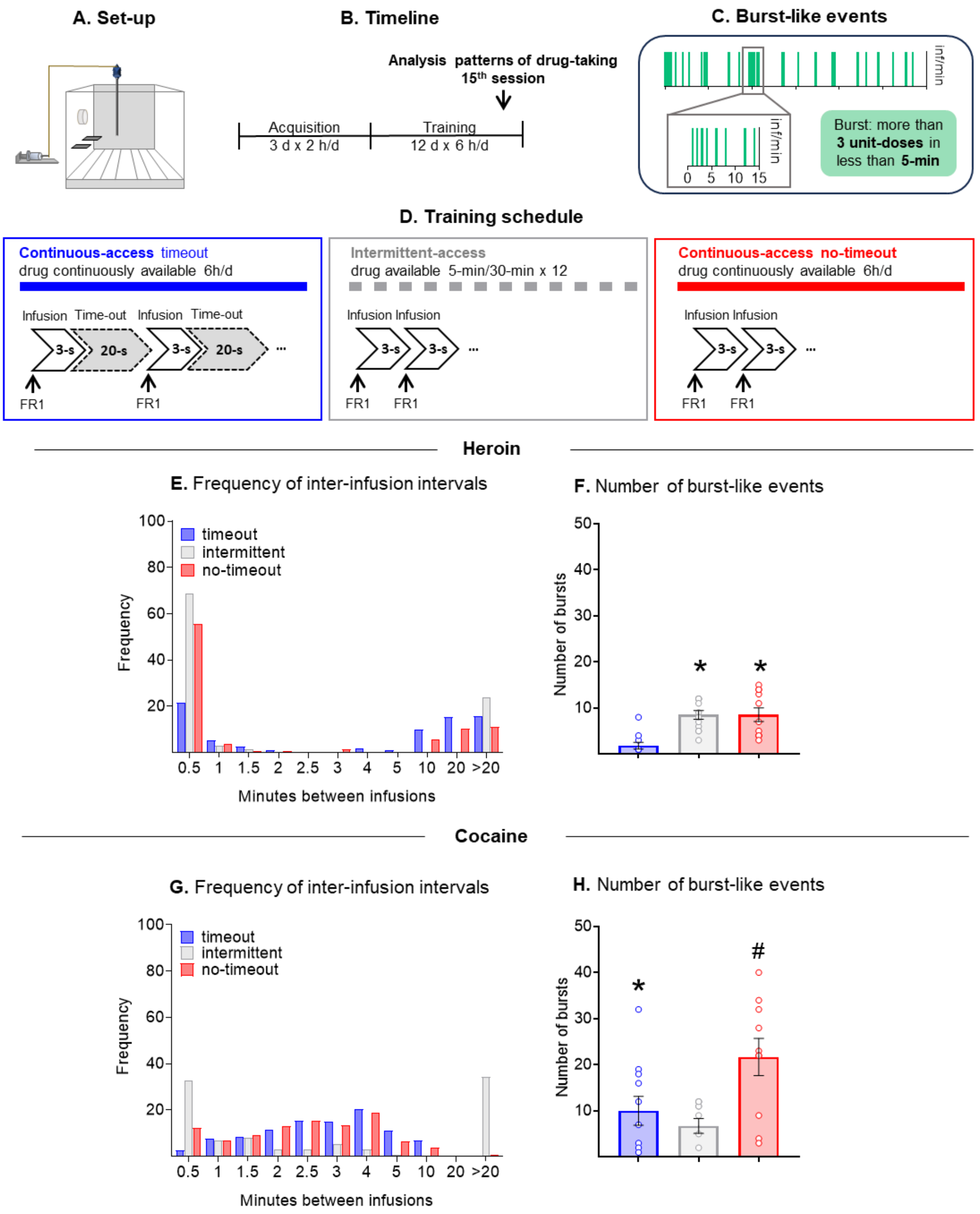
A. Set-up
B. Timeline
Analysis patterns of drug-taking
15th session
Acquisition
3 d x 2 h/d
Training
12 d x 6 h/d
C. Burst-like events
inf/min
0 5 10 15
Burst: more than 3 unit-doses in less than 5-min
D. Training schedule
Continuous-access timeout
drug continuously available 6h/d
Infusion Time-out Infusion Time-out
3-s 20-s 3-s 20-s ...
FR1 FR1
Intermittent-access
drug available 5-min/30-min x 12
Infusion Infusion
3-s 3-s ...
FR1 FR1
Continuous-access no-timeout
drug continuously available 6h/d
Infusion Infusion
3-s 3-s ...
FR1 FR1
Heroin
E. Frequency of inter-infusion intervals
timeout
intermittent
no-timeout
Frequency
Minutes between infusions
0.5 1 1.5 2 2.5 3 4 5 10 20 >20
F. Number of burst-like events
Number of bursts
Cocaine
G. Frequency of inter-infusion intervals
timeout
intermittent
no-timeout
Frequency
Minutes between infusions
0.5 1 1.5 2 2.5 3 4 5 10 20 >20
H. Number of burst-like events
Number of bursts

*Fig S4. Estimated brain concentrations of heroin and its active metabolites in representative rats trained under continuous-access timeout, intermittent-access or continuous-access no-timeout to heroin in the last session of drug self-administration training.*

(**A**) Experimental set-up. (**B**) Experimental timeline. (**C**) Training schedule. For each line: estimated drug brain concentrations (left) of (**D**) heroin, (**E**) 6-MAM and (**F**) morphine in representative rats (infusions: long-access = 30, intermittent-access = 55, continuous-access = 86); individual data of mean drug brain peak concentrations (right) of (**D**) heroin, (**E**) 6-MAM and (**F**) morphine. Note the scales are adapted to the levels of each compound. * Different from long-access, $p < 0.05$; † Different from continuous-access timeout and no-timeout, $p < 0.05$; # Different from continuous-access timeout and intermittent-access, $p < 0.05$. (n=37).

Figure S4.

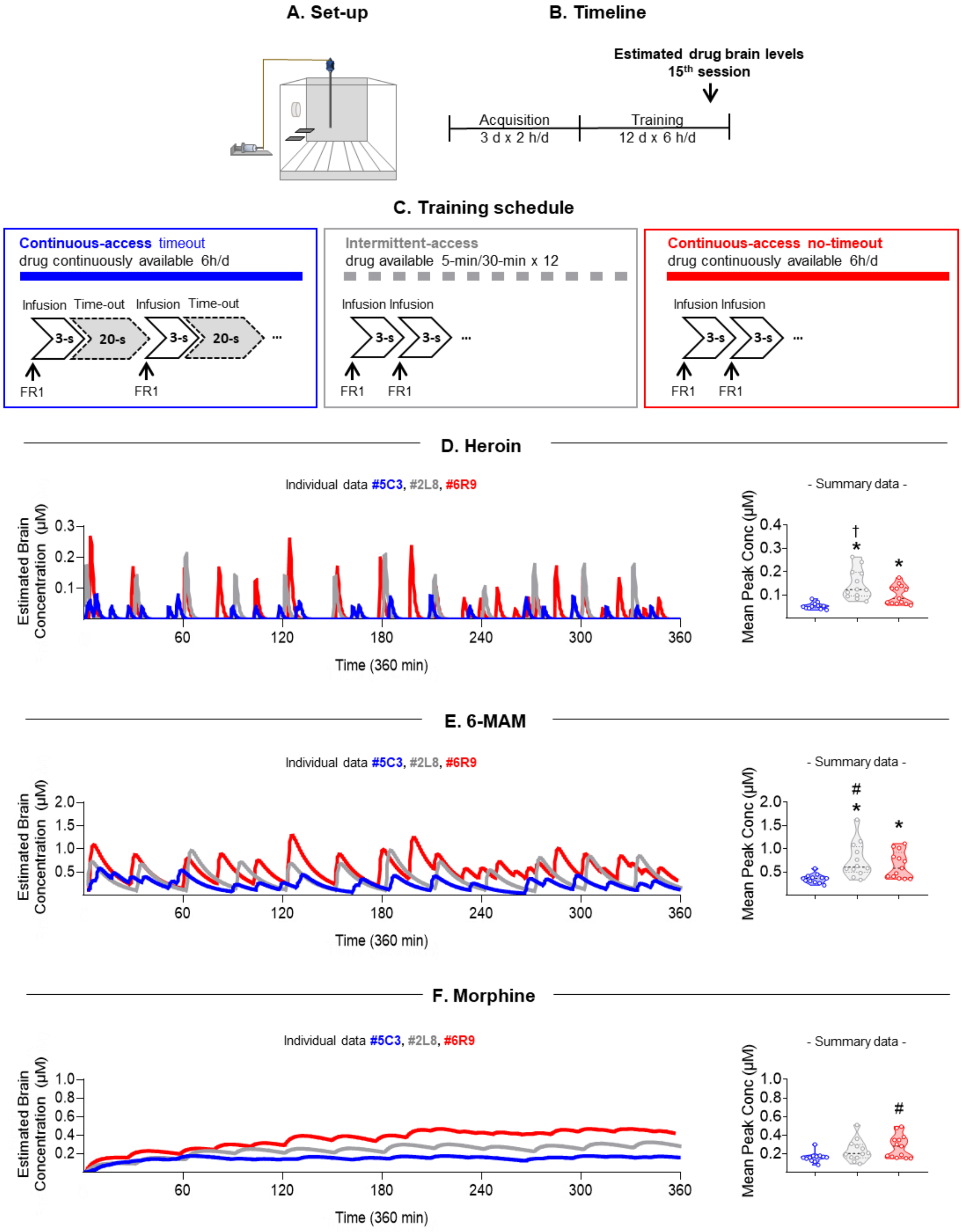
A. Set-up
B. Timeline
Estimated drug brain levels
15th session
Acquisition
3 d x 2 h/d
Training
12 d x 6 h/d
C. Training schedule
Continuous-access timeout
drug continuously available 6h/d
Infusion Time-out Infusion Time-out
3-s 20-s 3-s 20-s ...
FR1 FR1
Intermittent-access
drug available 5-min/30-min x 12
Infusion Infusion
3-s 3-s ...
FR1 FR1
Continuous-access no-timeout
drug continuously available 6h/d
Infusion Infusion
3-s 3-s ...
FR1 FR1
D. Heroin
Individual data #5C3, #2L8, #6R9
Estimated Brain Concentration (µM)
Time (360 min)
- Summary data -
Mean Peak Conc (µM)
E. 6-MAM
Individual data #5C3, #2L8, #6R9
Estimated Brain Concentration (µM)
Time (360 min)
- Summary data -
Mean Peak Conc (µM)
F. Morphine
Individual data #5C3, #2L8, #6R9
Estimated Brain Concentration (µM)
Time (360 min)
- Summary data -
Mean Peak Conc (µM)

*Figure S5. Estimated brain concentrations of heroin and its active metabolites in representative rats self-administering heroin without timeout and a simulated rat self-administering heroin with timeout.*

(**A**) Training schedule. (**B**) Pattern of drug-taking. (**C**) Estimated brain concentrations of heroin. (**D**) Estimated brain concentrations of 6-monoacthylmorphine (6-MAM). (**E**) Estimated brain concentrations of morphine.

**Figure S5.**

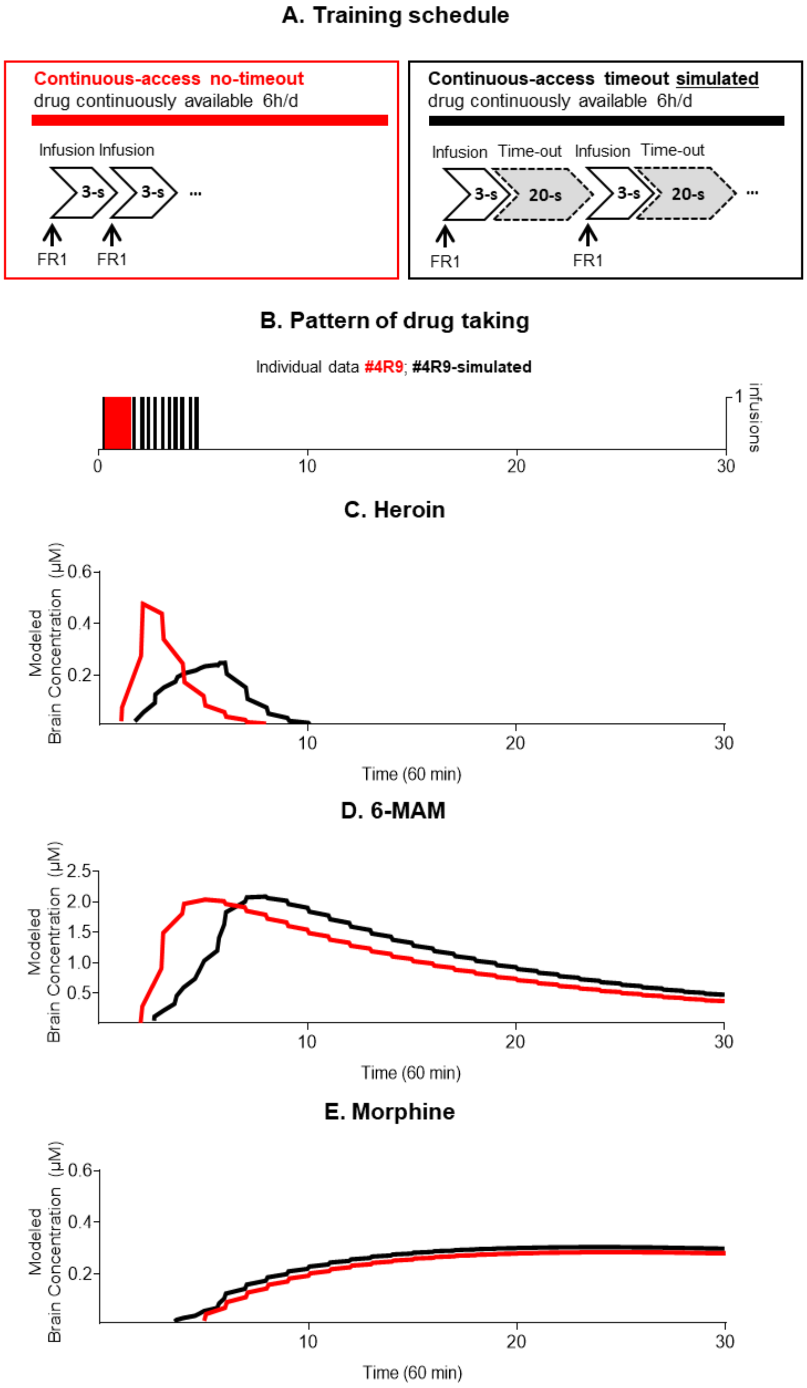
A. Training schedule
Continuous-access no-timeout
drug continuously available 6h/d
Infusion Infusion
3-s
3-s
FR1
FR1
Continuous-access timeout simulated
drug continuously available 6h/d
Infusion Time-out Infusion Time-out
3-s
20-s
3-s
20-s
FR1
FR1
B. Pattern of drug taking
Individual data #4R9; #4R9-simulated
1 infusions
0
10
20
30
C. Heroin
Modeled Brain Concentration (µM)
0.6
0.4
0.2
10
20
30
Time (60 min)
D. 6-MAM
Modeled Brain Concentration (µM)
2.5
2.0
1.5
1.0
0.5
10
20
30
Time (60 min)
E. Morphine
Modeled Brain Concentration (µM)
0.6
0.4
0.2
10
20
30
Time (60 min)

*Figure S6. Behavioral repertoire assessment in the first hour of the last session of drug self-administration training.*

(**A**) Experimental set-up. (**B**) Experimental timeline. Individual data of total time (s) of observed (**C and I**) stupor, (**D and J**) walking, (**E and K**) chewing, (**F and L**) grooming, (**G and M**) sniffing, (**H and N**) hand licking. * Different from long-access, $p < 0.05$; # Different from long-access and intermittent-access, $p < 0.05$; † Different from continuous-access timeout and no-timeout, $p < 0.05$. (heroin: n=31; cocaine: n=31).

**Figure S6.**

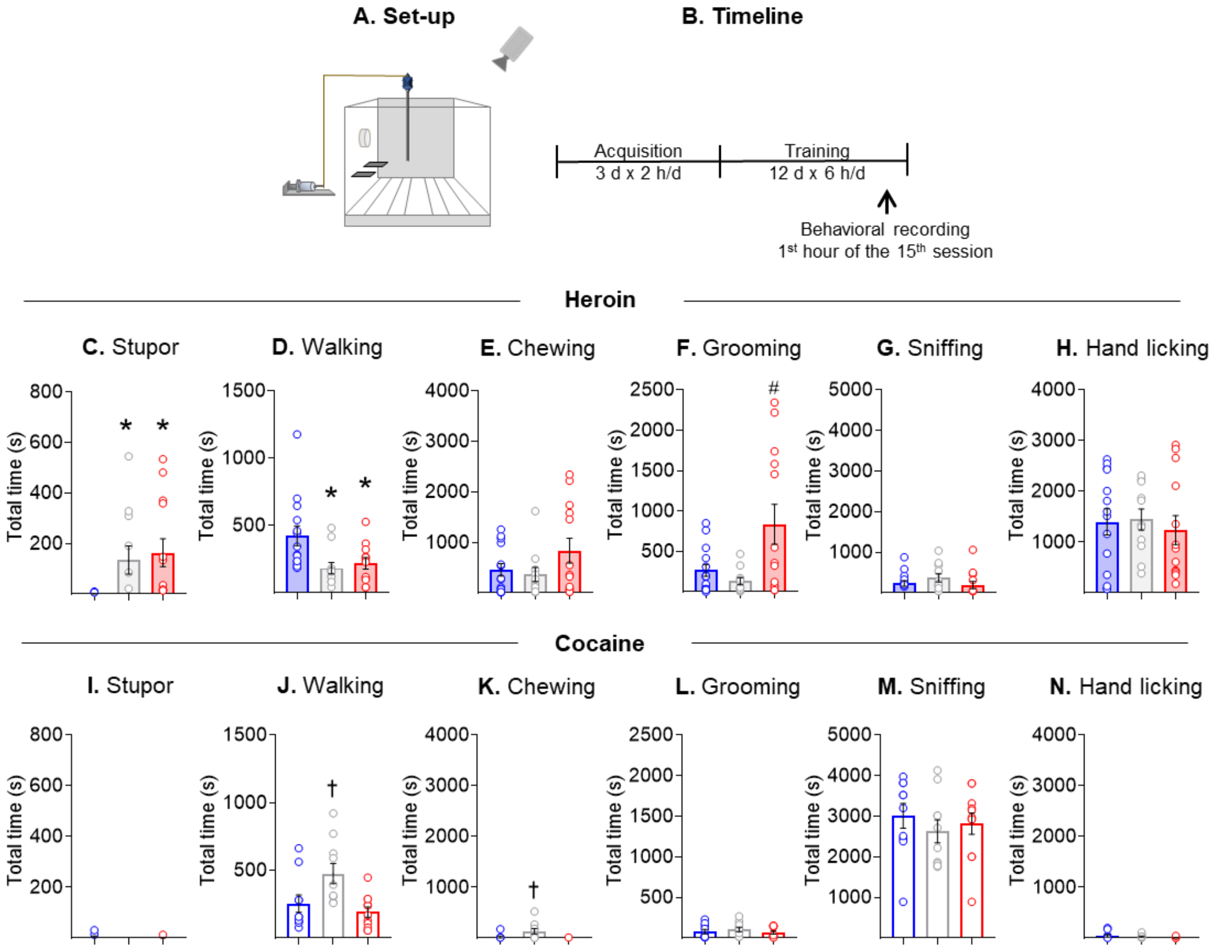
A. Set-up
B. Timeline
Acquisition
3 d x 2 h/d
Training
12 d x 6 h/d
Behavioral recording
1st hour of the 15th session
Heroin
C. Stupor
D. Walking
E. Chewing
F. Grooming
G. Sniffing
H. Hand licking
Cocaine
I. Stupor
J. Walking
K. Chewing
L. Grooming
M. Sniffing
N. Hand licking
Total time (s)

*Figure S7. Effects of procedural manipulations on drug/social preference: sampling and discrete choice.*

(**A**) Experimental set-up. (**B**) Experimental timeline. (**C and E**) Sampling and choice schedule. (**D**) Discrete choice. Heroin vs social interaction with incremental time of access to the social partner. Mean ± SEM preference score across all the access time conditions. (**F**) Discrete choice. Cocaine vs social interaction with incremental time of access to cocaine. Mean ± SEM preference score across all the access time conditions (heroin: n = 15; cocaine: n = 10).

**Figure S7.**

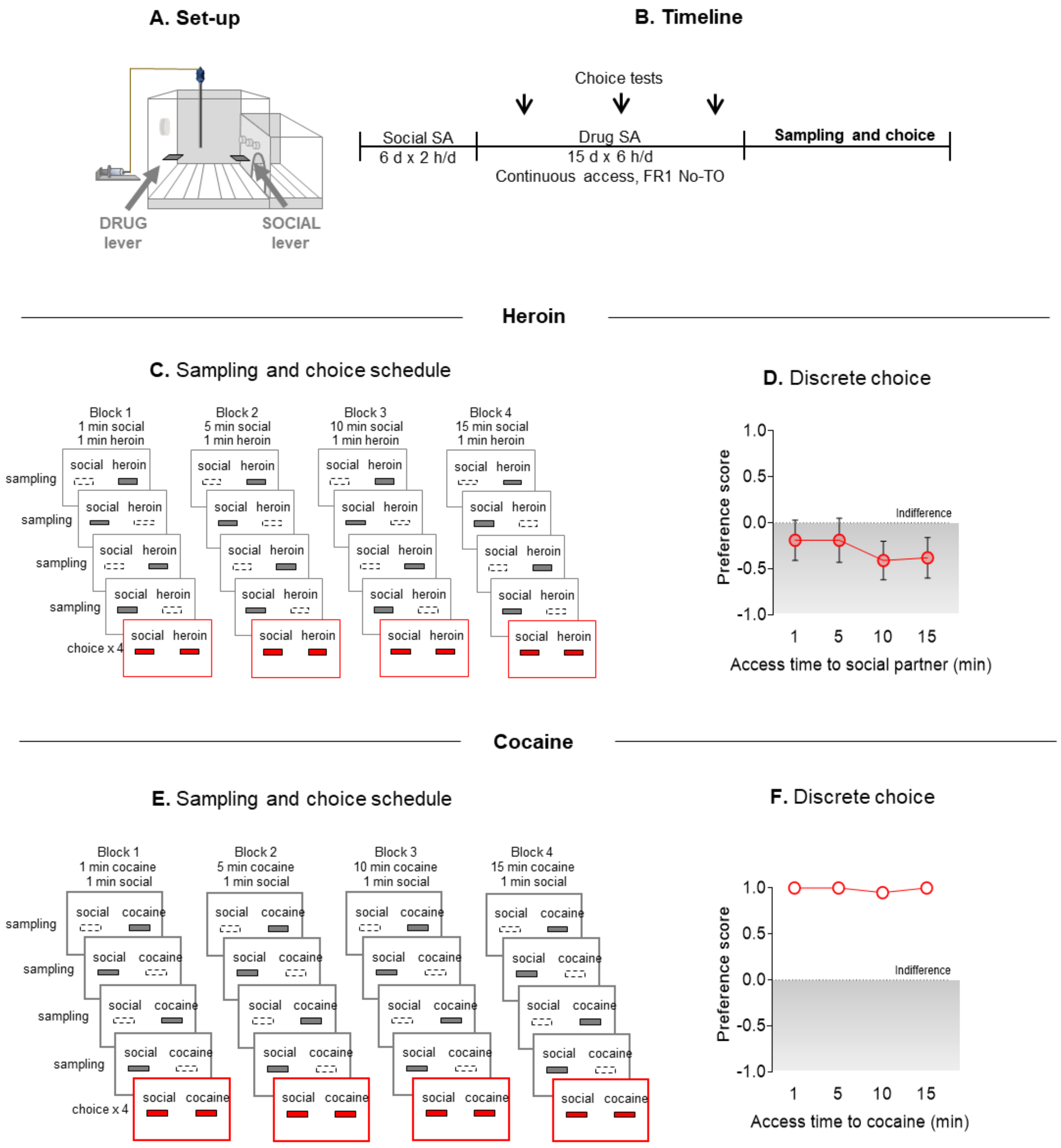
A. Set-up
DRUG lever
SOCIAL lever
B. Timeline
Choice tests
Social SA
6 d x 2 h/d
Drug SA
15 d x 6 h/d
Continuous access, FR1 No-TO
Sampling and choice
Heroin
C. Sampling and choice schedule
Block 1
1 min social
1 min heroin
Block 2
5 min social
1 min heroin
Block 3
10 min social
1 min heroin
Block 4
15 min social
1 min heroin
sampling
sampling
sampling
sampling
choice x 4
social heroin
D. Discrete choice
Preference score
1.0
0.5
0.0
-0.5
-1.0
Indifference
1
5
10
15
Access time to social partner (min)
Cocaine
E. Sampling and choice schedule
Block 1
1 min cocaine
1 min social
Block 2
5 min cocaine
1 min social
Block 3
10 min cocaine
1 min social
Block 4
15 min cocaine
1 min social
sampling
sampling
sampling
sampling
choice x 4
social cocaine
F. Discrete choice
Preference score
1.0
0.5
0.0
-0.5
-1.0
Indifference
1
5
10
15
Access time to cocaine (min)

*Figure S8. Discrete choice between drugs and social interaction after 30 days of abstinence.*

(**A**) Experimental set-up. (**B**) Experimental timeline. (**C**) Preference score heroin. (**D**) Preference score cocaine. (heroin: n =16 , cocaine n = 19).

**Figure S8**

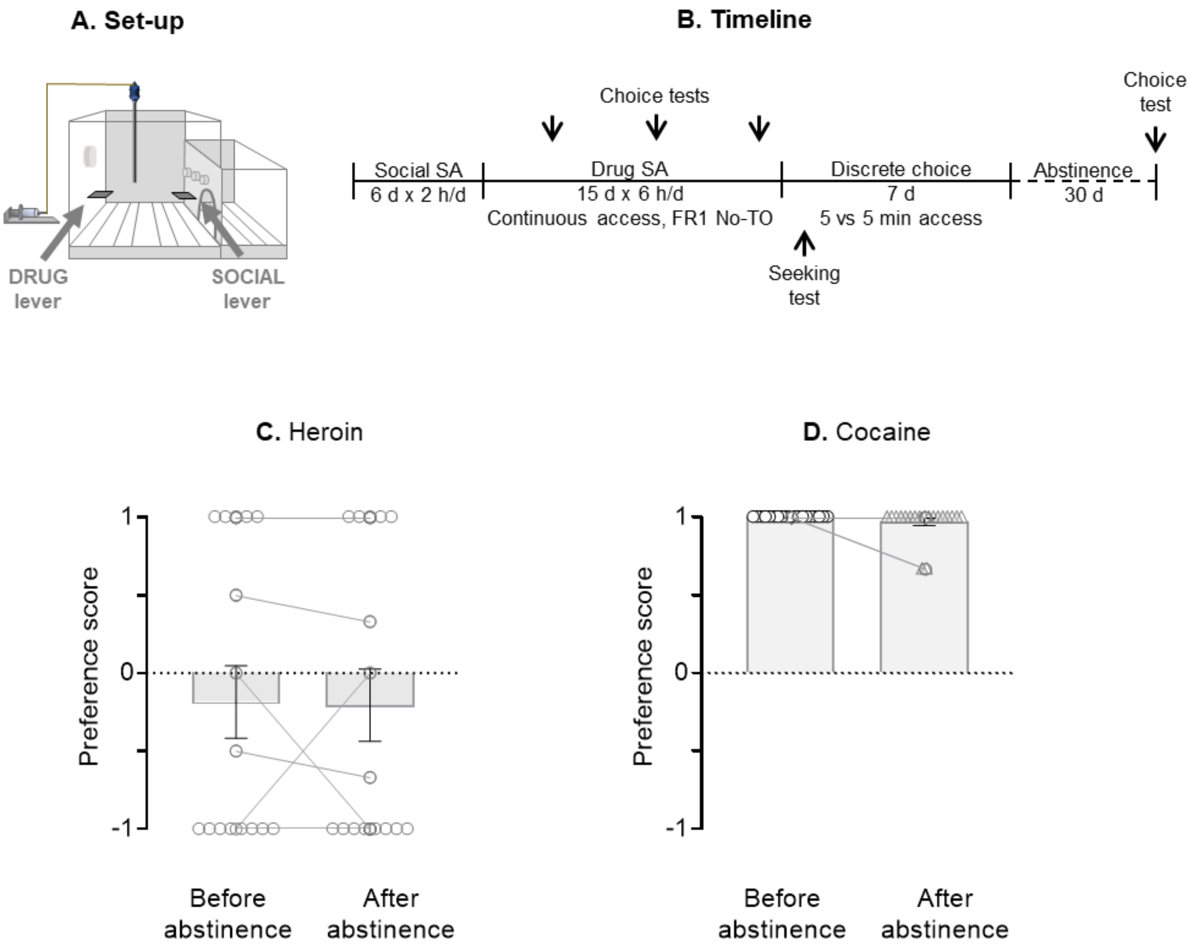
A. Set-up
DRUG lever
SOCIAL lever
B. Timeline
Choice tests
Choice test
Social SA
6 d x 2 h/d
Drug SA
15 d x 6 h/d
Continuous access, FR1 No-TO
Discrete choice
7 d
5 vs 5 min access
Abstinence
30 d
Seeking test
C. Heroin
D. Cocaine
Preference score
1
0
-1
Before abstinence
After abstinence

*Figure S9. Cluster analysis of rats tested on the '5 vs 5 min' choice procedure.*

(**A**) Experimental set-up. (**B**) Experimental timeline. (**C**) Features used for cluster analysis. (**D and E**) Dendrogram cluster analysis.

**Figure S9.**

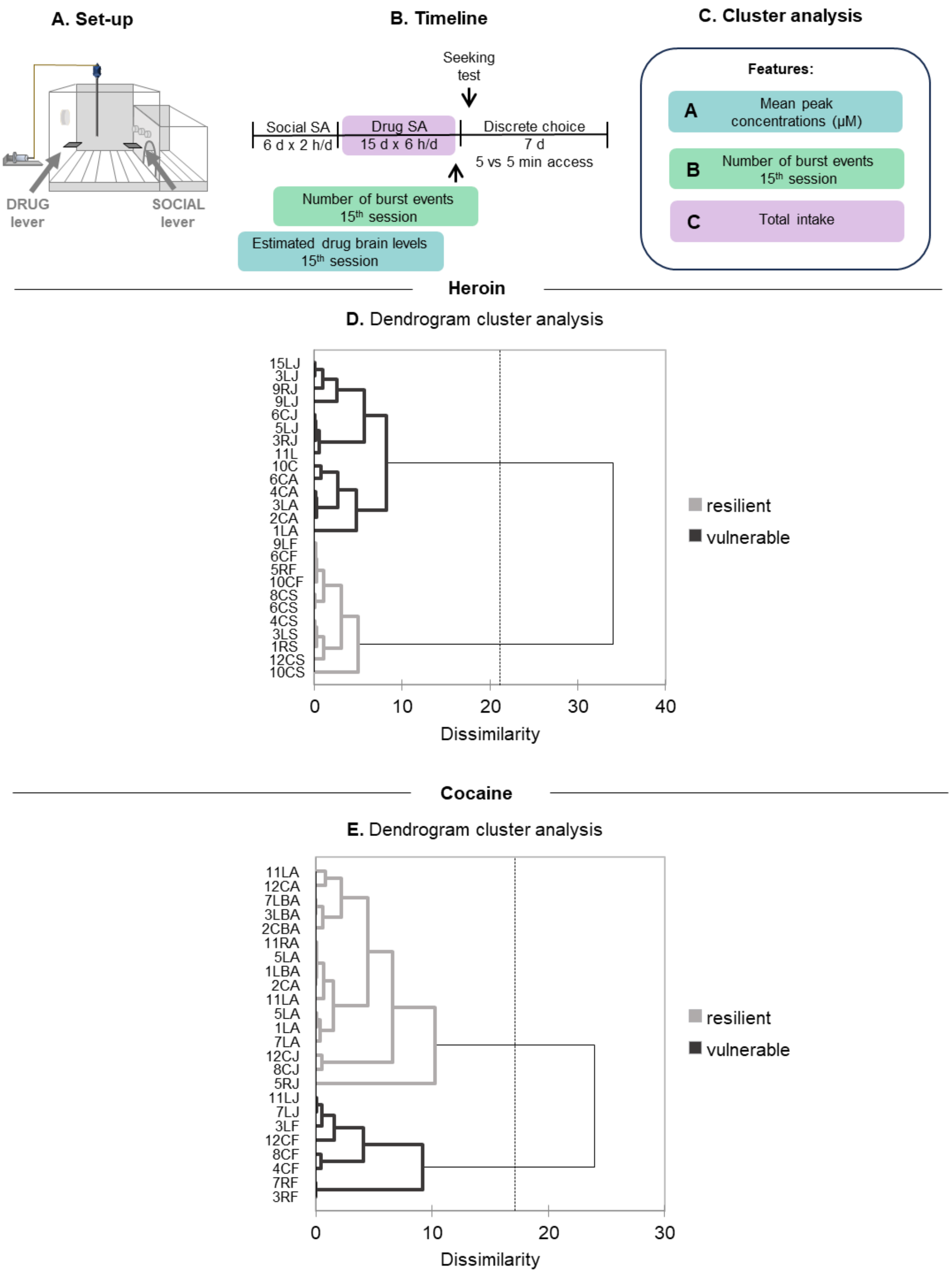
A. Set-up
DRUG lever
SOCIAL lever
B. Timeline
Seeking test
Social SA 6 d x 2 h/d
Drug SA 15 d x 6 h/d
Discrete choice 7 d 5 vs 5 min access
Number of burst events 15th session
Estimated drug brain levels 15th session
C. Cluster analysis
Features:
A Mean peak concentrations (μM)
B Number of burst events 15th session
C Total intake
Heroin
D. Dendrogram cluster analysis
15LJ 3LJ 9RJ 9LJ 6CJ 5LJ 3RJ 11L 10C 6CA 4CA 3LA 2CA 1LA 9LF 6CF 5RF 10CF 8CS 6CS 4CS 3LS 1RS 12CS 10CS
0 10 20 30 40
Dissimilarity
resilient
vulnerable
Cocaine
E. Dendrogram cluster analysis
11LA 12CA 7LBA 3LBA 2CBA 11RA 5LA 1LBA 2CA 11LA 5LA 1LA 7LA 12CJ 8CJ 5RJ 11LJ 7LJ 3LF 12CF 8CF 4CF 7RF 3RF
0 10 20 30
Dissimilarity
resilient
vulnerable

*Figure S10. Correlations and interindividual variability in drug-related behaviors in rats clusterized as vulnerable and resilient.*

(**A**) Experimental set-up. (**B**) Experimental timeline. (**C**) Features used for cluster analysis. (**D**) Severity x Seeking. Correlation between Severity z-score and Seeking. (**E**) Seeking. Drug seeking in resilient and vulnerable rats. (**F**) Severity x Choice. Correlation between Severity z-score and Choice. (**F**) Preference score. Social preference in resilient and vulnerable rats (heroin n=25, cocaine n=24)

**Figure S10.**

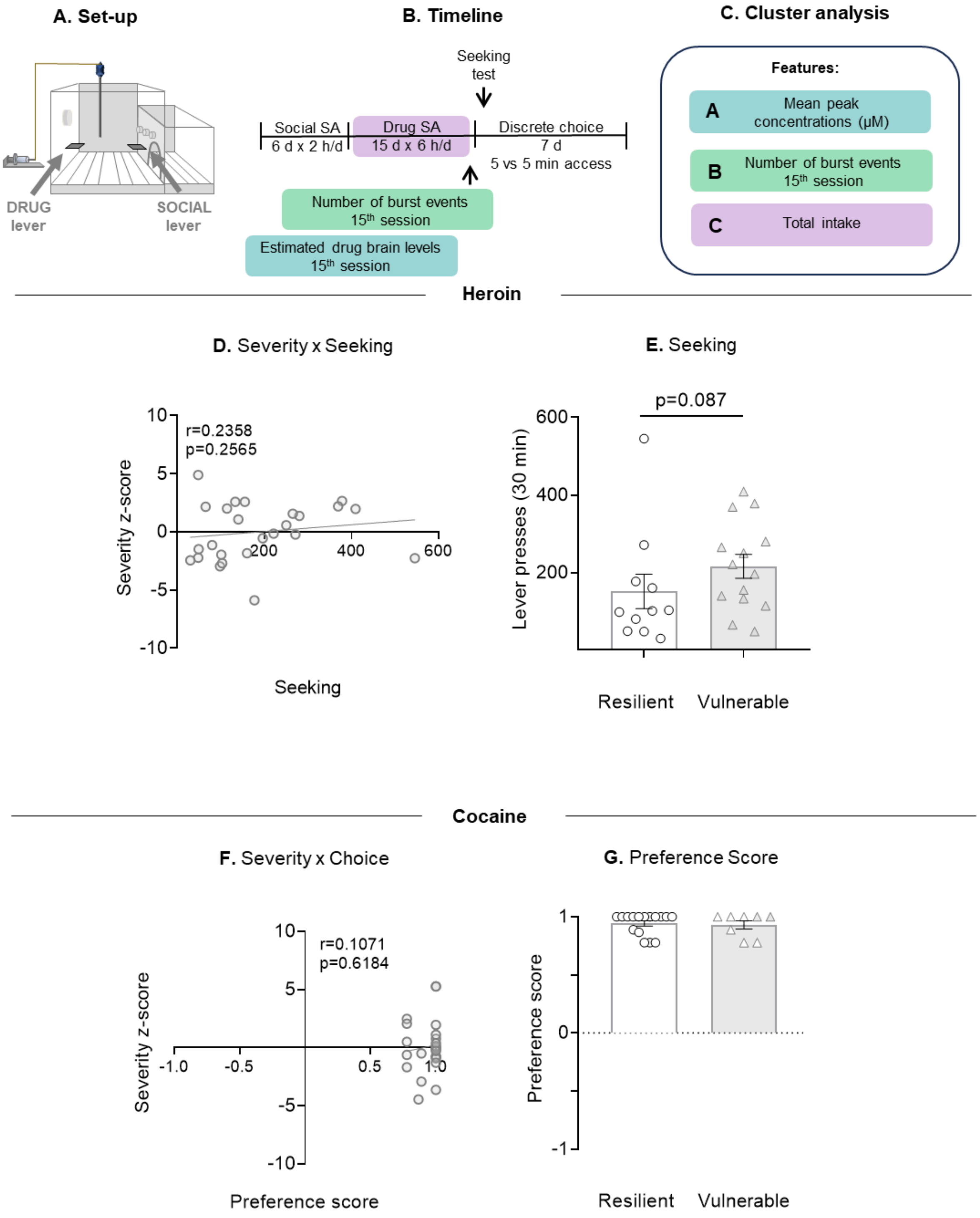
A. Set-up
DRUG lever
SOCIAL lever
B. Timeline
Seeking test
Social SA
6 d x 2 h/d
Drug SA
15 d x 6 h/d
Discrete choice
7 d
5 vs 5 min access
Number of burst events 15th session
Estimated drug brain levels 15th session
C. Cluster analysis
Features:
A Mean peak concentrations (µM)
B Number of burst events 15th session
C Total intake
Heroin
D. Severity x Seeking
r=0.2358
p=0.2565
Severity z-score
Seeking
E. Seeking
p=0.087
Lever presses (30 min)
Resilient
Vulnerable
Cocaine
F. Severity x Choice
r=0.1071
p=0.6184
Severity z-score
Preference score
G. Preference Score
Preference score
Resilient
Vulnerable

**Table S1.**

Pharmacokinetics parameters used in Gottås et al. [4] to fit the concentrations of heroin, 6-monoacetylmorphine (6-MAM), and morphine in the brain extracellular fluid after intravenous administration of 3 μmol (1.3 mg) heroin in the rat (Boix, Andersen & Mørland, 2013). These same parameters were applied to the FitMultiMicroExtravascular model of the software program Kinetica v.5.1 (Thermo Fisher Scientific Inc., Waltham, MA, USA) to simulate the brain concentrations considering the times of the single unit-doses during the 10th self-administration training session.

[1] One compartment extravascular model

[2] Two compartments extravascular model

[3] Three compartments extravascular model

Ka: Absorption rate constant from the injection site. Lag: Time taken to appear in the brain following administration. Kel: Elimination rate constant from the brain. Volume: Volume of distribution. KXY: Transfer rate constant between compartments X and Y (1 = measured compartment)

| **Parameter** | **Ka** | **lag** | **Volume** | **Kel** | **K12** | **K21** | **K13** | **K31** |
|---|---|---|---|---|---|---|---|---|
| Unit | $min^{-1}$ | min | L | $min^{-1}$ | $min^{-1}$ | $min^{-1}$ | $min^{-1}$ | $min^{-1}$ |
| Heroin[1] | 0.749072 | 0.757626 | 0.401859 | 2.18171 | | | | |
| 6-MAM[2] | 1.00001 | 1.52153 | 0.417121 | 0.0695501 | 0.00802414 | 0.032289 | | |
| Morphine[3] | 0.050732 | 1.22008 | 0.873335 | 0.0418547 | 0.114092 | 0.115256 | 0.0182552 | 0.012605 |

**Table S2.**

Pharmacokinetics parameters extracted from Pan et al. [5] used to fit the concentrations of cocaine in the brain extracellular fluid after intravenous administration of 7.5 mg/kg cocaine in the rat. These same parameters were applied to the FitMultiMicroExtravascular model of the software program Kinetica v.5.1 (Thermo Fisher Scientific Inc., Waltham, MA, USA) to simulate the brain concentrations taking into account the times of the single unit-doses during the last self-administration training session.

[1] Extravascular two compartments model without lag

Ka: Absorption rate constant from injection site. Kel: Elimination rate constant from brain. Volume: Volume of distribution. KXY: Transfer rate constant between compartment X and Y (1 = measured compartment).

| **Parameter** | **Ka** | **lag** | **Volume** | **Kel** | **K12** | **K21** | **K13** | **K31** |
|---|---|---|---|---|---|---|---|---|
| Unit | $min^{-1}$ | min | L | $min^{-1}$ | $min^{-1}$ | $min^{-1}$ | $min^{-1}$ | $min^{-1}$ |
| Cocaine[1] | 0.586706 | | 0.122182 | 0.090295 | 0.067599 | 0.047336 | | |

**Table S3.**

Statistical analysis (SPSS GLM repeated-measures module; R GLMM Generalized Linear Mixed-Effects Model; GraphPad Prism; XLSTAT).

| Figure | Data | Primary statistic | Post-hoc test | Comparison | p-value | F/t/r/z statistic |
|---|---|---|---|---|---|---|
| **Figure 1D** | Heroin training Intake | Generalized Linear Mixed-Effects Model | | | p=0.07 | |
| **Figure 1D** | Heroin training Total intake | Two-tailed, paired t-test | | Timeout vs No-Timeout | **p=0.001*** | z=13.500 |
| **Figure 1E** | Heroin training Lever presses | Generalized Linear Mixed-Effects Model | | | **p = 0.021*** | |
| **Figure 1F** | Cocaine training Intake | Generalized Linear Mixed-Effects Model | | | p=0.75 | |
| **Figure 1F** | Cocaine training Total intake | Two-tailed, paired t-test | | Timeout vs No-Timeout | | |
| **Figure 1G** | Cocaine training Lever presses | Generalized Linear Mixed-Effects Model | | | p=0.85 | |
| **Figure 2D** | Heroin seeking test | Two-tailed, paired t-test | | Timeout vs No-Timeout | **p=0.001*** | z=-4.011 |
| **not shown** | Heroin training Intake Access assignment (choice or PR test) | One-way RM ANOVA | | Access x Session interaction | p=0.344 | $F_{13,351}$=1.129 |
| | | | | Main effect of Access | **p=0.001*** | $F_{1,27}$=0.014 |
| | | | | Main effect of Session | p=0.908 | $F_{13,351}$=12.832 |
| **not shown** | Preference score | One-way RM ANOVA | | Main effect of Session | **p=0.001*** | $F_{8,144}$=4.685 |
| **Figure 2E** | Preference score | Two-tailed, paired t-test | | Timeout vs No-Timeout | **p=0.001*** | $T_{18}$=-4.693212 |
| **Figure 2F** | Progressive ratio (Total infusions) | Two-tailed, paired t-test | | Timeout vs No-Timeout | **p=0.012*** | z=-2.524 |
| **Figure 2F** | Progressive ratio (Final ratio) | Two-tailed, paired t-test | | Timeout vs No-Timeout | p=0.2717 | $T_{10}$=-1.163 |
| **Figure 2F** | Progressive ratio (Cumulative lever presses) | Two-tailed, paired t-test | | Timeout vs No-Timeout | **p=0.047*** | z=1.988 |
| **Figure 2G** | Cocaine seeking test | Two-tailed, paired t-test | | Timeout vs No-Timeout | **p=0.025*** | z=2.244 |

| | | | | | | |
|---|---|---|---|---|---|---|
| **not shown** | Cocaine training Intake Access assignment (choice or PR test) | One-way RM ANOVA | | Access x Time interaction | p=0.510 | $F_{13,338}$=0.801 |
| | | | | Main effect of Access | **p=0.001*** | $F_{13,338}$=27.088 |
| | | | | Main effect of Session | p=0.908 | $F_{1,26}$=0.585 |
| **not shown** | Preference score | 1-way RM ANOVA | | Main effect of Time | **p=0.001*** | $F_{8,144}$=4.685 |
| **Figure 2H** | Preference score | Two-tailed, paired t-test | | Timeout vs No-Timeout | **p=0.010*** | T16=-2.928311 |
| **Figure 2I** | Progressive ratio (Total infusions) | Two-tailed, paired t-test | | Timeout vs No-Timeout | **p=0.041*** | z=-2.047 |
| **Figure 2I** | Progressive ratio (Final ratio) | Two-tailed, paired t-test | | Timeout vs No-Timeout | **p=0.041*** | z=-2.047 |
| **Figure 2I** | Progressive ratio (Cumulative lever presses) | Two-tailed, paired t-test | | Timeout vs No-Timeout | **p=0.047*** | z=1.988 |
| **Figure 3D** | Heroin acquisition Intake | Two-way RM ANOVA | | Access x Session interaction | p=0.088 | $F_{4,68}$=2.612 |
| | | | | Main effect of Access | p=0.174 | $F_{2,34}$=1.843 |
| | | | | Main effect of Session | **p=0.001*** | $F_{2,68}$=15.975 |
| | | | PLSD | Continuous timeout vs Intermittent | p=0.137 | |
| | | | PLSD | Continuous timeout vs Continuous no-timeout | p=0.822 | |
| | | | PLSD | Intermittent vs Continuous no-timeout | p=0.084 | |
| **not shown** | Heroin acquisition Lever pressing | Two-way RM ANOVA | | Access x Session interaction | p=0.131 | $F_{4,68}$=1.967 |
| | | | | Main effect of Access | p=0.066 | $F_{2,34}$=2.955 |
| | | | | Main effect of Session | **p=0.001*** | $F_{2,68}$=8.427520 |
| | | | PLSD | Continuous timeout vs Intermittent | p=0.452070 | |
| | | | PLSD | Continuous timeout vs Continuous no-timeout | p=0.119196 | |
| | | | PLSD | Intermittent vs Continuous no-timeout | p=0.023374 | |
| **Figure 3D** | Heroin training Intake | Two-way RM ANOVA | | Access x Session interaction | **p=0.002*** | $F_{22,374}$=2.159322 |
| | | | | Main effect of Access | **p=0.001*** | $F_{2,34}$=4.573282 |

| | | | | | | |
|---|---|---|---|---|---|---|
| | | | | Main effect of Session | **p=0.001*** | $F_{11,374}$=17.061538 |
| | | | PLSD | Continuous timeout vs Intermittent | p=0.014970 | |
| | | | PLSD | Continuous timeout vs Continuous | p=0.011052 | |
| | | | PLSD | Intermittent vs Continuous no-timeout | p=0.941453 | |
| **not shown** | Heroin training Lever pressing | Two-way RM ANOVA | | Access x Session interaction | p=0.551659 | $F_{22,374}$=0.932557 |
| | | | | Main effect of Access | p=0.352560 | $F_{2,34}$=1.075164 |
| | | | | Main effect of Session | **p=0.001*** | $F_{22,374}$=9.885 |
| | | | PLSD | Continuous timeout vs Intermittent | p=0.168837 | |
| | | | PLSD | Continuous timeout vs Continuous no-timeout | p=0.722435 | |
| | | | PLSD | Intermittent vs Continuous no-timeout | p=0.289704 | |
| **3F** | Heroin seeking test AD1 | One-way ANOVA | | Main effect of Access | **p=0.003*** | $F_{2,34}$=6.904106 |
| | | | PLSD | Continuous timeout vs Intermittent | **p=0.001*** | |
| | | | PLSD | Continuous timeout vs Continuous no-timeout | **p=0.008*** | |
| | | | PLSD | Intermittent vs Continuous no-timeout | p=0.450 | |
| **Figure 3F** | Heroin seeking test AD1 (Time course) | Two-way RM ANOVA | | Access x Time interaction | p=0.380 | $F_{4,68}$=1.066165 |
| | | | | Main effect of Access | **p=0.003*** | $F_{2,34}$=6.904106 |
| | | | | Main effect of Time | **p=0.001*** | $F_{2,68}$=18.424894 |
| | | | PLSD | Continuous timeout vs Intermittent | **p=0.001*** | |
| | | | PLSD | Continuous timeout vs Continuous no-timeout | **p=0.008*** | |
| | | | PLSD | Intermittent vs Continuous no-timeout | p=0.450 | |
| **Figure 3G** | Cocaine acquisition Intake | Two-way RM ANOVA | | Access x Session interaction | p=0.298 | $F_{4,76}$=1.248610 |

| | | | | | | |
|---|---|---|---|---|---|---|
| | | | | Main effect of Access | p=0.989 | $F_{2,38}$=0.010858 |
| | | | | Main effect of Session | **p=0.044*** | $F_{2,76}$=3.244245 |
| | | | PLSD | Continuous timeout vs Intermittent | p=0.968043 | |
| | | | PLSD | Continuous timeout vs Continuous no-timeout | p=0.885801 | |
| | | | PLSD | Intermittent vs Continuous no-timeout | p=0.930287 | |
| **not shown** | Cocaine acquisition Lever pressing | Two-way RM ANOVA | | Access x Session interaction | p=0.746 | $F_{4,76}$=0.486377 |
| | | | | Main effect of Access | p=0.494 | $F_{2,38}$=0.719322 |
| | | | | Main effect of Session | p=0.971 | $F_{2,76}$=0.029248 |
| | | | PLSD | Continuous timeout vs Intermittent | p=0.429579 | |
| | | | PLSD | Continuous timeout vs Continuous no-timeout | p=0.253236 | |
| | | | PLSD | Intermittent vs Continuous no-timeout | p=0.822640 | |
| **Figure 3H** | Cocaine training Intake | Two-way RM ANOVA | | Access x Session interaction | **p=0.001*** | $F_{22,418}$=4.706878 |
| | | | | Main effect of Access | **p=0.001*** | $F_{2,38}$=47.056141 |
| | | | | Main effect of Session | **p=0.001*** | $F_{11,418}$=47.606853 |
| | | | PLSD | Continuous timeout vs Intermittent | **p=0.001*** | |
| | | | PLSD | Continuous timeout vs Continuous no-timeout | **p=0.026*** | |
| | | | PLSD | Intermittent vs Continuous no-timeout | **p=0.001*** | |
| **not shown** | Cocaine training Lever pressing | Two-way RM ANOVA | | Access x Session interaction | **p=0.004*** | $F_{22,418}$=2.057190 |
| | | | | Main effect of Access | **p=0.001*** | $F_{2,38}$=35.004707 |
| | | | | Main effect of Session | **p=0.001*** | $F_{11,418}$=23.446173 |
| | | | PLSD | Continuous timeout vs Intermittent | **p=0.001*** | |
| | | | PLSD | Continuous timeout vs | p=0.248799 | |

| | | | | | | |
|---|---|---|---|---|---|---|
| | | | | Continuous no-timeout | | |
| | | | PLSD | Intermittent vs Continuous no-timeout | **p=0.001*** | |
| **Figure 3I** | Cocaine seeking test AD1 | One-way ANOVA | | Main effect of Access | **p=0.001*** | $H_{2,37}$=13.06 |
| | | | PLSD | Continuous timeout vs Intermittent | **p=0.003*** | |
| | | | PLSD | Continuous timeout vs Continuous no-timeout | **p=0.015*** | |
| | | | PLSD | Intermittent vs Continuous no-timeout | p=0.211759 | |
| **Figure 3I** | Cocaine seeking test AD1 (Time course) | Two-way RM ANOVA | | Access x Time interaction | **p=0.002*** | $F_{2,76}$=4.778906 |
| | | | | Main effect of Access | **p=0.003*** | $F_{2,38}$=6.705851 |
| | | | | Main effect of Time | **p=0.001*** | $F_{2,76}$=71.675902 |
| | | | PLSD | Continuous timeout vs Intermittent | **p=0.001*** | |
| | | | PLSD | Continuous timeout vs Continuous | **p=0.018*** | |
| | | | PLSD | Intermittent vs Continuous no-timeout | p=0.176919 | |
| **not shown** | Heroin intake last self-administration session | One-way ANOVA | | Main effect of Access | **p=0.012*** | $F_{2,34}$=5.102462 |
| | | | PLSD | Continuous timeout vs Intermittent | p=0.071686 | |
| | | | PLSD | Continuous timeout vs Continuous no-timeout | **p=0.003*** | |
| | | | PLSD | Intermittent vs Continuous no-timeout | p=0.206866 | |
| **Figure S3F** | Number of bursts-like events (heroin) | One-way ANOVA | | Main effect of Access | **p=0.001*** | $F_{2,30}$=11.164 |
| | | | PLSD | Continuous timeout vs Intermittent | **p=0.001*** | |
| | | | PLSD | Continuous timeout vs Continuous no-timeout | **p=0.003*** | |
| | | | PLSD | Intermittent vs Continuous no-timeout | p=0.977 | |
| **Figure S3H** | Number of bursts-like | One-way ANOVA | | Main effect of Access | **p=0.004*** | $F_{2,31}$=6.599 |

| | | | | | | |
|---|---|---|---|---|---|---|
| | events (cocaine) | | PLSD | Continuous timeout vs Intermittent | p=0.442 | |
| | | | PLSD | Continuous timeout vs Continuous no-timeout | **p=0.011*** | |
| | | | PLSD | Intermittent vs Continuous no-timeout | **p=0.002*** | |
| **not shown** | Mean heroin infusions per peak | One-way ANOVA | | Main effect of Access | **p=0.001*** | $F_{2,34}$=11.432818 |
| | | | PLSD | Continuous timeout vs Intermittent | **p=0.001*** | |
| | | | PLSD | Continuous timeout vs Continuous no-timeout | **p=0.013*** | |
| | | | PLSD | Intermittent vs Continuous no-timeout | **p=0.031*** | |
| **Figure 4E** | Mean brain concentration of heroin per peak | One-way ANOVA | | Main effect of Access | **p=0.001*** | $H_{3,37}$=21.63 |
| | | | Dunn | Continuous timeout vs Intermittent | **p=0.001*** | |
| | | | Dunn | Continuous timeout vs Continuous no-timeout | **p=0.005*** | |
| | | | Dunn | Intermittent vs Continuous no-timeout | p=0.425 | |
| **not shown** | Mean heroin peaks slope | One-way ANOVA | | Main effect of Access | **p=0.001*** | $F_{2,34}$=12.829708 |
| | | | PLSD | Continuous timeout vs Intermittent | **p=0.001*** | |
| | | | PLSD | Continuous timeout vs Continuous no-timeout | **p=0.014*** | |
| | | | PLSD | Intermittent vs Continuous no-timeout | **p=0.014*** | |
| **Figure 4F** | Mean brain concentration of heroin | One-way ANOVA | | Main effect of Access | **p=0.029*** | $H_{2,34}$=7.076 |
| | | | Dunn | Continuous timeout vs Intermittent | p=0.1219 | |
| | | | Dunn | Continuous timeout vs Continuous no-timeout | **p=0.008*** | |
| | | | Dunn | Intermittent vs Continuous no-timeout | p=0.2833 | |
| **not shown** | Cocaine intake last self- | One-way ANOVA | | Main effect of Access | **p=0.001*** | $F_{2,38}$=65.662886 |

| | | | | | | |
|---|---|---|---|---|---|---|
| | administration session | | PLSD | Continuous timeout vs Intermittent | **p=0.001*** | |
| | | | PLSD | Continuous timeout vs Continuous no-timeout | p=0.052 | |
| | | | PLSD | Intermittent vs Continuous no-timeout | **p=0.001*** | |
| **not shown** | Mean cocaine infusions per peak | One-way ANOVA | | Main effect of Access | **p=0.011*** | $F_{2,38}$=5.037645 |
| | | | PLSD | Continuous timeout vs Intermittent | **p=0.003*** | |
| | | | PLSD | Continuous timeout vs Continuous no-timeout | p=0.241857 | |
| | | | PLSD | Intermittent vs Continuous | **p=0.038*** | |
| **Figure 4H** | Mean brain concentration of cocaine per peak | One-way ANOVA | | Main effect of Access | **p=0.001*** | $H_{3,38}$=17.16 |
| | | | Dunn | Continuous timeout vs Intermittent | **p=0.021*** | |
| | | | Dunn | Continuous timeout vs Continuous no-timeout | p=0.3361 | |
| | | | Dunn | Intermittent vs Continuous no-timeout | **p=0.001*** | |
| **not shown** | Mean cocaine peaks slope | One-way ANOVA | | Main effect of Access | **p=0.001*** | $F_{2,38}$=7.833116 |
| | | | PLSD | Continuous timeout vs Intermittent | **p=0.001*** | |
| | | | PLSD | Continuous timeout vs Continuous no-timeout | p=0.491967 | |
| | | | PLSD | Intermittent vs Continuous no-timeout | **p=0.003*** | |
| **Figure 4I** | Mean brain concentration of cocaine | One-way ANOVA | | Main effect of Access | **p=0.001*** | $H_{2,38}$=39.0636 |
| | | | Dunn | Continuous timeout vs Intermittent | **p=0.001*** | |
| | | | Dunn | Continuous timeout vs Continuous no-timeout | p=0.4000 | |
| | | | Dunn | Intermittent vs Continuous no-timeout | **p=0.001*** | |
| **Figure S4E** | Mean brain concentration of | One-way ANOVA | | Main effect of Access | **p=0.004*** | $F_{2,34}$=6.576353 |

| | | | | | | |
|---|---|---|---|---|---|---|
| | 6-MAM per peak | | PLSD | Continuous timeout vs Intermittent | **p=0.001*** | |
| | | | PLSD | Continuous timeout vs Continuous no-timeout | **p=0.014*** | |
| | | | PLSD | Intermittent vs Continuous no-timeout | p=0.341533 | |
| **Figure S4F** | Mean brain concentration of morphine per peak | One-way ANOVA | | Main effect of Access | **p=0.027*** | $F_{2,34}$=4.027738 |
| | | | PLSD | Continuous timeout vs Intermittent | p=0.086302 | |
| | | | PLSD | Continuous timeout vs Continuous no-timeout | **p=0.008*** | |
| | | | PLSD | Intermittent vs Continuous no-timeout | p=0.319473 | |
| **Figure S1D** | Heroin seeking test AD21 | One-way ANOVA | | Main effect of Access | p=0.4306 | $H_{2,37}$=9.885 |
| | | | PLSD | Continuous timeout vs Intermittent | p=0.6291 | |
| | | | PLSD | Continuous timeout vs Continuous no-timeout | p=0.999 | |
| | | | PLSD | Intermittent vs Continuous no-timeout | p=0.999 | |
| **Figure S1D** | Heroin seeking test AD1-AD21 (incubation) | Two-way RM ANOVA | | Main effect of Access | **p=0.002*** | $F_{2,38}$=3.568233 |
| | | | | Main effect of Time | p=0.154903 | $F_{2,38}$=2.106263 |
| | | | | Access x Time interaction | **p=0.038*** | $F_{2,38}$=3.568233 |
| | | | PLSD | Continuous timeout vs Intermittent | **p=0.001*** | |
| | | | PLSD | Continuous timeout vs Continuous no-timeout | **p=0.042*** | |
| | | | PLSD | Intermittent vs Continuous no-timeout | p=0.054 | |
| **Figure S1E** | Heroin body weight | Two-way RM ANOVA | | Main effect of Access | p=0.050833 | $F_{2,34}$=3.256211 |
| | | | | Main effect of Session | **p=0.001*** | $F_{14,476}$=11.640652 |
| | | | | Access x Session interaction | p=0.111988 | $F_{28,476}$=1.978551 |

| | | | | | | |
|---|---|---|---|---|---|---|
| | | | PLSD | Continuous timeout vs Intermittent | **p=0.041*** | |
| | | | PLSD | Continuous timeout vs Continuous no-timeout | **p=0.028*** | |
| | | | PLSD | Intermittent vs Continuous no-timeout | p=0.902470 | |
| **Figure S1F** | Heroin Von Frey test (hyperalgesia) | Two-way RM ANOVA | | Main effect of Access | p=0.721673 | $F_{2,35}$=0.329241 |
| | | | | Main effect of Time | **p=0.001*** | $F_{3,105}$=64.784474 |
| | | | | Access x Time interaction | p=0.653393 | $F_{6,105}$=0.695964 |
| | | | PLSD | Continuous timeout vs Intermittent | p=0.467960 | |
| | | | PLSD | Continuous timeout vs Continuous no-timeout | p=0.954505 | |
| | | | PLSD | Intermittent vs Continuous no-timeout | p=0.502545 | |
| **Figure S1G** | Cocaine seeking test AD21 | One-way ANOVA | | Main effect of Access | **p=0.009*** | $F_{2,38}$=5.379362 |
| | | | PLSD | Continuous timeout vs Intermittent | **p=0.003*** | |
| | | | PLSD | Continuous timeout vs Continuous no-timeout | p=0.439134 | |
| | | | PLSD | Intermittent vs Continuous no-timeout | **p=0.015*** | |
| **Figure S1G** | Cocaine seeking test AD1-AD21 (incubation) | Two-way RM ANOVA | | Main effect of Access | **p=0.002*** | $F_{1,38}$=7.370770 |
| | | | | Main effect of Time | p=0.154903 | $F_{2,38}$=2.106263 |
| | | | | Access x Time interaction | **p=0.038*** | $F_{2,38}$=3.568233 |
| | | | PLSD | Continuous timeout vs Intermittent | **p=0.001*** | |
| | | | PLSD | Continuous timeout vs Continuous no-timeout | **p=0.042*** | |
| | | | PLSD | Intermittent vs Continuous no-timeout | p=0.054 | |
| **Figure S1H** | Cocaine body weight | Two-way RM ANOVA | | Main effect of Access | p=0.497680 | $F_{2,28}$=0.715480 |
| | | | | Main effect of Session | **p=0.001*** | $F_{14,392}$=177.701531 |

| | | | | | | |
|---|---|---|---|---|---|---|
| | | | | Access x Session interaction | **p=0.001*** | $F_{28,392}$=8.894700 |
| | | | PLSD | Continuous timeout vs Intermittent | p=0.670820 | |
| | | | PLSD | Continuous timeout vs Continuous no-timeout | p=0.245480 | |
| | | | PLSD | Intermittent vs Continuous no-timeout | p=0.465851 | |
| **Figure S6C** | Stupor | One-way ANOVA | | Main effect of Access | **p=0.022*** | $F_{2,35}$=4.272452 |
| | | | PLSD | Continuous timeout vs Intermittent | **p=0.038*** | |
| | | | PLSD | Continuous timeout vs Continuous no-timeout | **p=0.010*** | |
| | | | PLSD | Intermittent vs Continuous no-timeout | **p=0.650** | |
| **Figure S6D** | Walking | One-way ANOVA | | Main effect of Access | **p=0.008*** | $F_{2,35}$=5.561852 |
| | | | PLSD | Continuous timeout vs Intermittent | **p=0.005*** | |
| | | | PLSD | Continuous timeout vs Continuous no-timeout | **p=0.010*** | |
| | | | PLSD | Intermittent vs Continuous no-timeout | p=0.711706 | |
| **Figure S6E** | Chewing | One-way ANOVA | | Main effect of Access | p=0.168282 | $F_{2,35}$=1.876017 |
| | | | PLSD | Continuous timeout vs Intermittent | p=0.723563 | |
| | | | PLSD | Continuous timeout vs Continuous no-timeout | p=0.135962 | |
| | | | PLSD | Intermittent vs Continuous no-timeout | p=0.082847 | |
| **Figure S6F** | Grooming | One-way ANOVA | | Main effect of Access | **p=0.007*** | $F_{2,35}$=5.706851 |
| | | | PLSD | Continuous timeout vs Intermittent | p=0.549023 | |
| | | | PLSD | Continuous timeout vs Continuous no-timeout | **p=0.011*** | |

| | | | | | | |
|---|---|---|---|---|---|---|
| | | | PLSD | Intermittent vs Continuous no-timeout | **p=0.003*** | |
| **Figure S6G** | Sniffing | One-way ANOVA | | Main effect of Access | p=0.350569 | $F_{2,35}$=1.080226 |
| | | | PLSD | Continuous timeout vs Intermittent | p=0.277315 | |
| | | | PLSD | Continuous timeout vs Continuous no-timeout | p=0.725470 | |
| | | | PLSD | Intermittent vs Continuous no-timeout | p=0.164989 | |
| **Figure S6H** | Hand licking | One-way ANOVA | | Main effect of Access | p=0.837110 | $F_{2,35}$=0.178706 |
| | | | PLSD | Continuous timeout vs Intermittent | p=0.893258 | |
| | | | PLSD | Continuous timeout vs Continuous no-timeout | p=0.652065 | |
| | | | PLSD | Intermittent vs Continuous no-timeout | p=0.578697 | |
| **Figure S6I** | Stupor | One-way ANOVA | | Main effect of Access | p=0.125447 | $F_{2,28}$=1.336917 |
| | | | PLSD | Continuous timeout vs Intermittent | p=0.251169 | |
| | | | PLSD | Continuous timeout vs Continuous no-timeout | p=0.659784 | |
| | | | PLSD | Intermittent vs Continuous no-timeout | p=0.578697 | |
| **Figure S6J** | Walking | One-way ANOVA | | Main effect of Access | **p=0.005*** | $F_{2,28}$=6.316419 |
| | | | PLSD | Continuous timeout vs Intermittent | **p=0.015*** | |
| | | | PLSD | Continuous timeout vs Continuous no-timeout | p=0.452044 | |
| | | | PLSD | Intermittent vs Continuous no-timeout | **p=0.002*** | |
| **Figure S6K** | Chewing | One-way ANOVA | | Main effect of Access | **p=0.011*** | $F_{2,28}$=5.375668 |
| | | | PLSD | Continuous timeout vs Intermittent | **p=0.016*** | |
| | | | PLSD | Continuous timeout vs Continuous no-timeout | p=0.676351 | |

| | | | | | | |
|---|---|---|---|---|---|---|
| | | | PLSD | Intermittent vs Continuous no-timeout | **p=0.005*** | |
| **Figure S6L** | Grooming | One-way ANOVA | | Main effect of Access | p=0.525433 | $F_{2,28}$=0.658553 |
| | | | PLSD | Continuous timeout vs Intermittent | p=0.436744 | |
| | | | PLSD | Continuous timeout vs Continuous no-timeout | p=0.757179 | |
| | | | PLSD | Intermittent vs Continuous no-timeout | p=0.272315 | |
| **Figure S6M** | Sniffing | One-way ANOVA | | Main effect of Access | p=0.639362 | $F_{2,28}$=0.454506 |
| | | | PLSD | Continuous timeout vs Intermittent | p=0.348536 | |
| | | | PLSD | Continuous timeout vs Continuous no-timeout | p=0.628988 | |
| | | | PLSD | Intermittent vs Continuous no-timeout | p=0.629811 | |
| **Figure S6N** | Hand licking | One-way ANOVA | | Main effect of Access | p=0.117956 | $F_{2,28}$=2.323206 |
| | | | PLSD | Continuous timeout vs Intermittent | p=0.244506 | |
| | | | PLSD | Continuous timeout vs Continuous no-timeout | p=0.051472 | |
| | | | PLSD | Intermittent vs Continuous no-timeout | p=0.326871 | |
| **Figure 5D** | Social self-administration | Two-way RM ANOVA | | Choice x Session interaction | p=0.898 | $F_{5,155}$=0.028257 |
| | | | | Main effect of Choice | p=0.867598 | $F_{1,31}$=0.028257 |
| | | | | Main effect of Session | **p=0.001*** | $F_{5,155}$=81.815445 |
| **Figure 5D** | Heroin self-administration | Two-way RM ANOVA | | Access x Session interaction | p=0.109025 | $F_{14,434}$=2.041447 |
| | | | | Main effect of Choice | p=0.748158 | $F_{1,31}$=0.104938 |
| | | | | Main effect of Session | **p=0.001*** | $F_{14,434}$=20.532306 |
| **Figure 5E** | Heroin seeking | Two-tailed, unpaired t-test | | 1:1 vs 5:5 Choice | p=0.563 | $T_{31}$=0.584 |
| **Figure 5F** | Preference score | Two-way RM ANOVA | | Choice x Session interaction | p=0.226284 | $F_{6,186}$=1.376076 |

| | | | | | | |
|---|---|---|---|---|---|---|
| | | | | Main effect of Choice | **p=0.002*** | $F_{1,31}$=11.250930 |
| | | | | Main effect of Session | p=0.990098 | $F_{6,186}$=0.143614 |
| **Figure 5F** | Preference score | Two-tailed, unpaired t-test | | 1:1 vs 5:5 Choice | **p=0.001*** | $T_{24.229}$=5.971613 |
| **Figure 5F** | Preference score (Choice 1 vs 1) | Two-tailed, paired t-test | | Social vs Heroin | **p=0.001*** | $T_{7}$=-81.992683 |
| **Figure 5F** | Preference score (Choice 5 vs 5) | Two-tailed, paired t-test | | Social vs Heroin | p=0.802304 | $T_{24}$=0.253156 |
| **Figure 5G** | Social self-administration | Two-way RM ANOVA | | Choice x Session interaction | p=0.509672 | $F_{5,155}$=0.859805 |
| | | | | Main effect of Choice | p=0.898299 | $F_{1,31}$=0.016606 |
| | | | | Main effect of Session | **p=0.001*** | $F_{5,155}$=105.119213 |
| **Figure 5G** | Cocaine self-administration | Two-way RM ANOVA | | Access x Session interaction | p=0.994890 | $F_{14,434}$=0.289621 |
| | | | | Main effect of Choice | p=0.937921 | $F_{1,31}$=0.006165 |
| | | | | Main effect of Session | **p=0.001*** | $F_{14,434}$=50.126369 |
| **Figure 5H** | Cocaine seeking | Two-tailed, unpaired t-test | | 1:1 vs 5:5 Choice | p=0.248795 | $T_{31}$=-1.175370 |
| **Figure 5I** | Preference score | Two-way RM ANOVA | | Choice x Session interaction | p=0.961218 | $F_{10,310}$=0.363913 |
| | | | | Main effect of Choice | p=0.628799 | $F_{1,31}$=0.238401 |
| | | | | Main effect of Session | **p=0.001*** | $F_{10,310}$=5.022495 |
| **Figure 5I** | Preference score | Two-tailed, unpaired t-test | | 1:1 vs 5:5 Choice | p=0.226799 | $F_{30.704}$=-1.233365 |
| **Figure 5I** | Preference score (Choice 1 vs 1) | Two-tailed, paired t-test | | Social vs Cocaine | **p=0.001*** | $T_{7}$=-63.561781 |
| **Figure 5I** | Preference score (Choice 5 vs 5) | Two-tailed, paired t-test | | Social vs Cocaine | **p=0.001*** | $T_{24}$=-72.883616 |
| **Figure S7D** | Preference score | One-way RM ANOVA | | Social vs Heroin | **p=0.046*** | $F_{3,45}$=3.617772 |
| **Figure S7F** | Preference score | One-way RM ANOVA | | Social vs Cocaine | p=0.406 | $F_{3,30}$=1.00000 |
| | Heroin (Choice 5 vs 5) | KMO | | | 0.768 | |
| | Heroin (Choice 5 vs 5) | Bartlett's Test | | | **p=0.001*** | $\chi^2$ =28.960 |
| | Cocaine (Choice 5 vs 5) | KMO | | | 0.600 | |
| | Cocaine (Choice 5 vs 5) | Bartlett's Test | | | **p=0.0489*** | $\chi^2$ = 18.376 |
| **not shown** | Factor 1 x Choice | Spearman Correlation | | | | r=-0.597 |

| | | | | | | |
|---|---|---|---|---|---|---|
| | Heroin (Choice 5 vs 5) | | | | | |
| **not shown** | Factor 1 x Seeking Cocaine (Choice 5 vs 5) | Spearman Correlation | | | | r=0.774 |
| **Figure 7D** | Severity x Choice | Spearman Correlation | | | **p=0.001*** | r=-0.6087 |
| **Figure 7E** | Sum Severity z-score | Two-tailed, unpaired t-test | | Resilient vs Vulnerable | **p=0.001*** | $H_{25}$=17.311 |
| **Figure 7F** | Preference score | Two-tailed, unpaired t-test | | Resilient vs Vulnerable | **p=0.007*** | $H_{25}$=7.311 |
| **Figure 7G** | Severity x Seeking | Spearman Correlation | | | **p=0.047*** | r=0.4078 |
| **Figure 7H** | Sum Severity z-score | Two-tailed, unpaired t-test | | Resilient vs Vulnerable | **p=0.001*** | $H_{24}$=10.244 |
| **Figure 7I** | Seeking | Two-tailed, unpaired t-test | | Resilient vs Vulnerable | **p=0.002*** | $H_{24}$=9.632 |
| **Figure S8C** | Preference score | Two-tailed, paired t-test | | Before and after abstinence | p=0.9704 | |
| **Figure S8D** | Preference score | Two-tailed, paired t-test | | Before and after abstinence | p=0.4865 | |
| **Figure S10D** | Severity x Seeking | Spearman Correlation | | | p=0.2565 | r=0.2358 |
| **Figure S10E** | Seeking | Two-tailed, unpaired t-test | | Resilient vs Vulnerable | p=0.085 | $H_{25}$=2.975 |
| **Figure S10F** | Severity x Choice | Spearman Correlation | | | p=0.6184 | r=0.1071 |
| **Figure S10G** | Preference score | Two-tailed, unpaired t-test | | Resilient vs Vulnerable | p=0.855 | $H_{24}$=0.034 |

**Table S4.** Contribution of the variables (%) after Varimax rotation for PCA on heroin (Choice 5 vs 5) dataset.

| | **Factor** | |
|---|---|---|
| **Variables** | **D1** | **D2** |
| **Z_Intake** | 31.381 | 0.016 |
| **Z_Peaks Conc** | 25.394 | 0.340 |
| **Z_Bursts Num** | 27.476 | 5.308 |
| **Z_Seeking** | 0.075 | 68.948 |
| **Z_Social Preference** | 15.674 | 25.388 |

**Table S5.** Squared cosines of the variables after Varimax rotation for PCA on heroin (Choice 5 vs 5) dataset. (Values in bold correspond for each variable to the factor for which the squared cosine is the largest).

| | **Factor** | |
|---|---|---|
| **Variables** | **D1** | **D2** |
| **Z_Intake** | **0.714** | 0.000 |
| **Z_Peaks Conc** | **0.578** | 0.004 |
| **Z_Bursts Num** | **0.625** | 0.069 |
| **Z_Seeking** | 0.002 | **0.899** |
| **Z_Social Preference** | **0.357** | 0.331 |

**Table S6.** Number of clusters for heroin (Choice 5 vs 5) dataset.

| **Number of clusters** | **2** | **3** | **4** | **5** |
|---|---|---|---|---|
| **Silhouette index** | 0.390 | 0.321 | 0.347 | 0.334 |
| **Hartigan index (H)** | 6.112 | 4.947 | 5.182 | 6.375 |
| **H(k-1) - H(k)** | **14.534** | 1.165 | -0.235 | -1.194 |
| **Calinski & Harabasz index** | 20.646 | 15.674 | 13.973 | 13.862 |

**Table S7.** Contribution of the variables (%) after Varimax rotation for PCA on cocaine (Choice 5 vs 5) dataset.

| | **Factor** | |
|---|---|---|
| **Variables** | **D1** | **D2** |
| **Z_Intake** | 8.330 | 1.809 |
| **Z_Peaks Conc** | 24.512 | 5.055 |
| **Z_Bursts Num** | 38.071 | 0.044 |
| **Z_Seeking** | 29.087 | 0.543 |
| **Z_Social Preference** | 0.001 | 92.549 |

**Table S8.** Squared cosines of the variables after Varimax rotation for PCA on cocaine (Choice 5 vs 5) dataset. (Values in bold correspond for each variable to the factor for which the squared cosine is the largest).

| | **Factor** | |
|---|---|---|
| **Variables** | **D1** | **D2** |
| **Z_Intake** | **0.172** | 0.019 |
| **Z_Peaks Conc** | **0.505** | 0.053 |
| **Z_Bursts Num** | **0.784** | 0.000 |
| **Z_Seeking** | **0.599** | 0.006 |
| **Z_Social Preference** | 0.000 | **0.961** |

**Table S9.** Number of clusters for cocaine  (Choice 5 vs 5) dataset.

| **Number of clusters** | **2** | **3** | **4** | **5** |
|---|---|---|---|---|
| **Silhouette index** | 0.317 | 0.318 | 0.330 | 0.348 |
| **Hartigan index (H)** | 6.338 | 7.455 | 6.927 | 5.849 |
| **H(k-1) - H(k)** | **5.608** | -1.117 | 0.529 | 1.077 |
| **Calinski & Harabasz index** | 11.946 | 10.592 | 11.717 | 13.123 |
